# Orchestrated long-distance gene activation by a ParB-like BisD-CTP DNA clamp in low-frequency transfer competence development in *Pseudomonas putida*

**DOI:** 10.1101/2024.09.16.613209

**Authors:** Hammam Antar, Nicolas Carraro, Stephan Gruber, Jan Roelof van der Meer

## Abstract

Integrative conjugative elements (ICEs) are mobile DNA that remain integrated within the host bacterial genome until they activate their excision and transfer to recipient cells via conjugation. ICE transfer is initiated in a small subpopulation of cells that undergo a hierarchical gene expression cascade leading to transfer competence formation. In this study, we demonstrate that transfer competence formation of the ICE*clc* element in stationary phase *Pseudomonas putida* cells is regulated by the coordinated activity of a two-component transcriptional activator, BisC and BisD. Chromatin immunoprecipitation of tagged BisC or BisD followed by high throughput sequencing showed that both proteins accumulate at similar sites around ICE*clc* transfer competence promoters in *P. putida* cells. Genetic dissection and single cell microscopy showed that BisD is a dual domain protein, with a C-terminal gene activator domain and an N-terminal ParB-like domain, forming dimeric DNA clamps that self-load at distant sites and reach target promoters after extensive one-dimensional DNA sliding. Expressed mCherry-BisD in *P. putida* ICE*clc* cells form discrete fluorescent foci, dependent on *parS-*like sequences on the ICE. This focus formation is similar as what is seen with canonical ParB proteins accumulating on chromosomal DNA, but in case of BisD corresponds to both chromosomal and excised ICE-molecules. Sliding of BisD over ∼50 kb of ICE-DNA is asymmetric and follows the direction of ICE gene transcription. This may help to establish a temporal order of activation of transfer competence formation, optimizing ICE transfer. Given that ICE*clc* is activated in stationary phase cells, we hypothesize that BisD is not involved in segregating excised ICE-DNA among daughter cells, but rather in actively directing ICE-DNA molecules towards the (multiple) conjugative systems that are produced in transfer competent cells. BisD thus serves as a twin function protein, integrating gene activation and DNA segregation functions.

## Introduction

ParABS systems are widely distributed on bacterial chromosomes, plasmids, and extrachromosomal prophages (Livny et al., 2007) and ensure the faithful segregation of genetic material during cell division by partitioning copies of the DNA within the bacterial cell (reviewed in (Jalal & Le, 2020)). The tripartite system consists of the ParB protein, acting as an adaptor connecting the *parS* DNA sequences to the ParA-ATPase motor proteins. ParB accumulates near *parS* sites at high concentrations generating distinct clusters within the cell. This accumulation involves ParB proteins binding CTP nucleotides and forming ParB DNA clamps, a reaction which is catalyzed by *parS* DNA (Jalal et al., 2020; Osorio-Valeriano et al., 2019; Soh et al., 2019). Subsequently, ParB DNA clamps are released from *parS* DNA and slide along the DNA double helix onto neighboring regions. CTP hydrolysis destabilizes the ParB clamp allowing for ParB-DNA dissociation, which then enables renewed ParB DNA clamping at *parS* DNA (Antar et al., 2021; Jalal et al., 2021; Osorio-Valeriano et al., 2021). Additional interactions between ParB and non-specific DNA sequences, as well as to other ParB dimers, are likely to contribute to the stabilization and further enlargement of the ParB cluster (Balaguer et al., 2021; Tisma et al., 2023; Tisma et al., 2022). Typically, ParB covers up to 10-15 kb of DNA flanking a *parS* site (Antar et al., 2021; Graham et al., 2014; Minnen et al., 2016). Variants of the ParABS systems have also been identified in mobile genetic elements such as integrative and conjugative elements or ICEs (Carraro, Sentchilo, et al., 2020; Reinhard et al., 2013), but their precise functions remain unclear. Here, we investigate a variant of ParABS encoded by the ICE*clc* element in *Pseudomonas putida* that contains a hybrid ParB protein, denoted as BisD, acting in conjunction with an activator protein BisC to initiate the expression of factors involved in ICE conjugative self-transfer (Carraro, Sentchilo, et al., 2020).

ICEs comprise groups of chromosomally integrated genetic elements with different evolutionary origins that transfer at low frequencies among related species (Delavat et al., 2017; Johnson & Grossman, 2015; Wozniak & Waldor, 2010). Transfer of the ICE is initiated by its excision from the chromosome through site-specific recombination, formation of a conjugative system, transfer of a single-stranded ICE-DNA molecule through the conjugative pore, replication in the new recipient cell and integration into the new host’s chromosome (summarized in (Johnson & Grossman, 2015)). ICEs have raised attention because they encode a large variety of functions affecting bacterial lifestyles, which as a result of their self-transfer capacity can lead to adaptation of new recipient strains under selective conditions. Examples of such cargo genes include resistances to heavy metals or to antimicrobial compounds, and catabolic pathways (Benigno et al., 2023; Carraro, Sentchilo, et al., 2020; Colombi et al., 2024; Colombi et al., 2017; Gaillard et al., 2008; Jones et al., 2021; Leon-Sampedro et al., 2016; Ryan et al., 2017; Zamarro et al., 2016).

A prominent example of adaptation is demonstrated in the ICE*clc* element discovered originally in *Pseudomonas knackmussii* B13 (Gaillard et al., 2006). The element was identified for its role in metabolizing 3-chlorobenzoate (3-CBA) by the *clcRABDE* operon genes that it carries (Miyazaki et al., 2015). These genes encode enzymes crucial for the degradation of <u>c</u>hloro<u>c</u>atechols (*clc*), formed as intermediates in 3-CBA metabolism. Conjugative transfer initiation of ICE*clc* occurs in a small percentage of cells (3–5%) in stationary phase, particularly after growth on 3-CBA (Minoia et al., 2008; Sentchilo et al., 2003). These cells activate a bistable switch, which starts a cascade of ICE*clc* gene expression, named the *transfer competence* program (Sulser et al., 2022), leading to a subpopulation of cells able to transfer the ICE. In contrast, a majority of cells keeps an inactive chromosomally integrated ICE*clc*. Cells with initiated transfer competence do not revert to the ICE-integrated state, and their cell division is eventually arrested, as a consequence of the ICE-encoded Shi and ParA proteins (Reinhard et al., 2013; Takano et al., 2019).

The activation of ICE*clc* transfer competence involves a regulatory hierarchy, passing step-by-step from the transcriptional factors MfsR, TciR (Pradervand et al., 2014), BisR, AlpA to BisDC (Carraro, Richard, et al., 2020). BisDC was thought to exist as a complex formed of BisD and BisC subunits, based on analogy to the *Escherichia coli* flagellar synthesis transcription factor FlhDC, even though only BisC has very low similarity to FlhC (Carraro, Richard, et al., 2020). Both components of BisDC are necessary for activation of its own expression, leading to a positive feedback loop that maintains bistable expression of the ICE transfer competence program (Carraro, Richard, et al., 2020). The transfer competence regulon involves at least eight further ICE*clc*-specific polycistronic units that are controlled by BisDC (Sulser et al., 2022), among which the formation of a type IV conjugation system (Daveri et al., 2023). Interestingly, however, part of BisD is homologous to ParB proteins and the *bisD* gene is located downstream of a *parA*-like gene on the ICE*clc* (Reinhard et al., 2013), suggesting that it is a hybrid protein with dual function in the ICE-transfer process: gene activation and ICE*clc* DNA segregation.

The major aim of this study was thus to elucidate the role of the BisD protein in ICE*clc* transfer and uncover its potential analogies to typical ParB functionalities. By gene deletions, expression of fluorescent-fusion proteins and epifluorescence microscopy, we show that BisD has two functional parts, and we delineate the BisD-parts essential for ICE gene activation. We show that BisD shares structural features with ParB proteins, loads, and accumulates around specific *parS*-like DNA sequences on ICE*clc*, herein referred to as the <u>bi</u>*<u>s</u>*tability regulatory <u>s</u>ites *bisS*. We purified the BisD protein and studied its binding to ribonucleotides as well as their hydrolysis in the presence of *bisS* DNA. We propose a CTP-dependent mechanism of BisD accumulation near *bisS* DNA. Additionally, we conducted chromatin immunoprecipitation to determine BisC and BisD distribution on chromosomal DNA, demonstrating close colocalization at known ICE-transfer competence-activated promoters in the core region of the ICE*clc* (Sulser et al., 2022). However, while BisC-binding is very specific and limited to those promoters sites, BisD accumulation showed a broad distribution covering essentially the entire ICE*clc* element with highest enrichment at *bisS*-proximal sites and the adjacent BisDC-activated promoters. With multiple target promoters being tens of kilobases distantly located from the nearest *bisS* site, we propose that the sliding of BisD DNA clamps along DNA is rate-limiting for distant BisDC target activation and may contribute to a stepwise activation of ICE*clc* genes. Overall, the results presented here highlights the prevalence of CTP-dependent loading and sliding, and provide novel insights and starting points for further investigations into ICE*clc* regulation and transfer.

## Results

### Structure prediction suggests BisD dimers to form a ParB-like DNA clamp

The BisD protein of ICE*clc* has a length of 550 amino acids making it significantly larger than ParB proteins. Sequence comparison by BLAST had previously suggested that BisD might be composed of two distinct parts: an N-terminal sequence (here designated as BisD_par_) with similarity to N-terminal sequences of ParB proteins, and a C-terminal domain with unclear function and no obvious sequence similarity (designated as BisD_act_ for activation domain) (Carraro, Richard, et al., 2020). AlphaFold3 (AF3) structure prediction of full-length BisD proteins reveals a tightly interacting clamp-like homodimer, with dimerization interfaces located at both the N- and the C-terminal parts of the protein (Fig. 1A). These parts are connected by an about 70-aa long apparently unstructured linker (Fig. 1A). The potential for dimerization can also be seen in AF3 predictions of isolated BisD_par_ and BisD_act_ (Fig. 1B). The folding of BisD_par_ strongly resembles the corresponding parts of canonical ParB proteins including a putative CTP-binding domain, with noticeable conservation of residues responsible for forming the CTP binding pocket in ParB (GxxR and ENxxR; in ParB proteins – G_92_NTR_95_, E_136_N_137_ELR_140_ in BisD; Fig. 1C) and a helix-turn-helix (HTH) domain, implying a capability of recognizing *parS*-like DNA sequences. A domain swap characteristic for ParB dimers (Fig. 1C upper panel) is also observed in the predicted BisD_par_ dimer structure, underscoring its similarities with ParB proteins (Antar et al., 2021). The unstructured flexible linker region in BisD is significantly longer than in ParB. In ParB, the linker is implicated in embracing chromosomal DNA after *parS* DNA loading, possibly implying that the extended linker in BisD supports particularly efficient DNA sliding or accommodates DNA-binding partner proteins in addition to a DNA double helix (Fig. 1C lower panel). BisD_act_ forms a tight homodimer in AF3 predictions comprising exclusively alpha-helical elements. The predicted structures do not show obvious similarity to known structures (including FlhDC; based on PDBeFold searches), and AF3 prediction together with BisC failed to reveal a robust interaction. Yet, BisC models confirmed low but detectable structural similarities of its C-terminal domain to FlhC (Fig. S1), although it lacks the zinc binding site implicated in DNA binding by FlhC and has an additional N-terminal dimerization domain that more distantly positions the putative DNA binding regions in the dimer. In summary, the AF3 predictions support the notion that BisD is a hybrid protein comprising two parts: one originating from a ParABS DNA partitioning system and another of less clear origin, possibly an FlhDC-like transcriptional activator.

**Figure 1:**
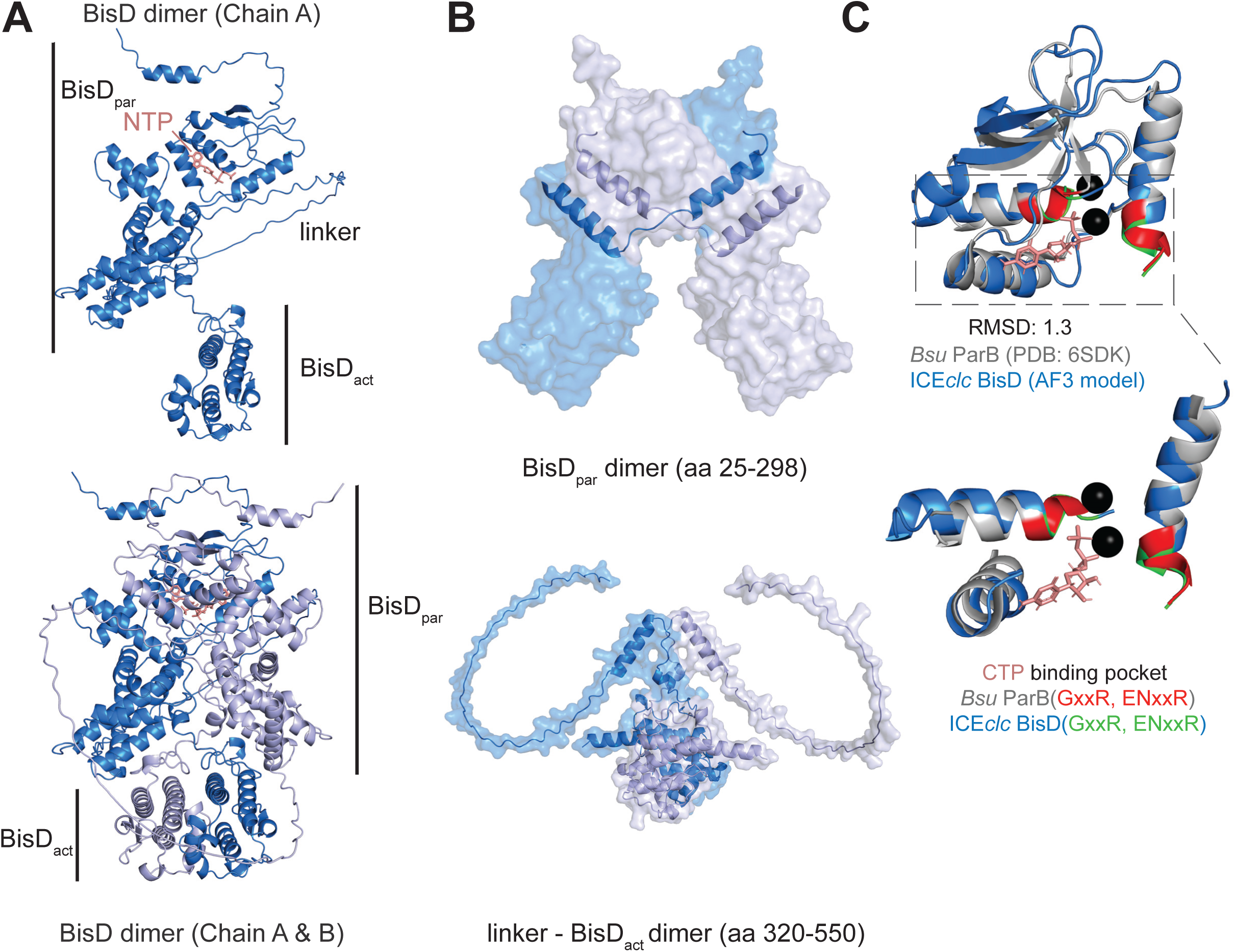
Structure prediction of ICEclc BisD rotein by AlphaFold3. *(A)* Full length BisD dimer model with the left panel showing only one chain, and the right panel showing both chains. The nucleotide (ATP in this case) is shown in salmon color. *(B)* Independent Alphafold3 prediction models for the BisD_par_ and BisD_act_ domains, with BisD_par_ folding exhibiting a domain swap characteristic of ParB proteins. *(C)* Superimposition of BisD_par_ domain (model in marine blue) and *Bacillus subtilis* ParB N-CDP (PDB:6SDK in gray) with the GxxR and the ENxxR motif of each protein highlighted (red for *B. subtilis* ParB and green for *P. putida* ICE*clc* BisD (RMSD= 1.3).

### The BisD DNA binding domain is needed for ICEclc transcription regulation

BisD and BisC have been implicated in transcriptional regulation of ICE*clc* transfer competence genes (Carraro, Richard, et al., 2020). To test the role of the potential BisD_act_ activator domain more specifically, a *P. putida* UWC derivative strain without ICE*clc* was constructed carrying a single copy chromosomally inserted dual fluorescent reporter. This reporter combines an *egfp* gene fused to a copy of the ICE*clc inrR* promoter, and an *echerry* gene fused to the *intB13* integrase promoter of the ICE (Fig. 2A). This strain with a plasmid containing either *bisD* or *bisC* under control of the *tac* promoter, did not show any appreciable levels of reporter fluorescence upon induction with IPTG (Fig. 2B), implying that expression of neither *bisC* nor *bisD* alone is sufficient for activation of these ICE-promoters. In contrast, simultaneous induction of *bisD* and *bisC* activated the reporters in all individual cells. Shortening *bisD* to a fragment encompassing only *bisD_act_* also gave rise to induction of both reporters in individual cells, but only when combined with *bisC* expression (Fig. 2B). This demonstrates that expression of BisD in combination with BisC is sufficient for activation of ICE promoters for transfer competence formation (Sulser et al., 2022). This activation does not require BisD_par_, at least when BisD is expressed from a plasmid-located IPTG-inducible *tac* promoter.

**Figure 2:**
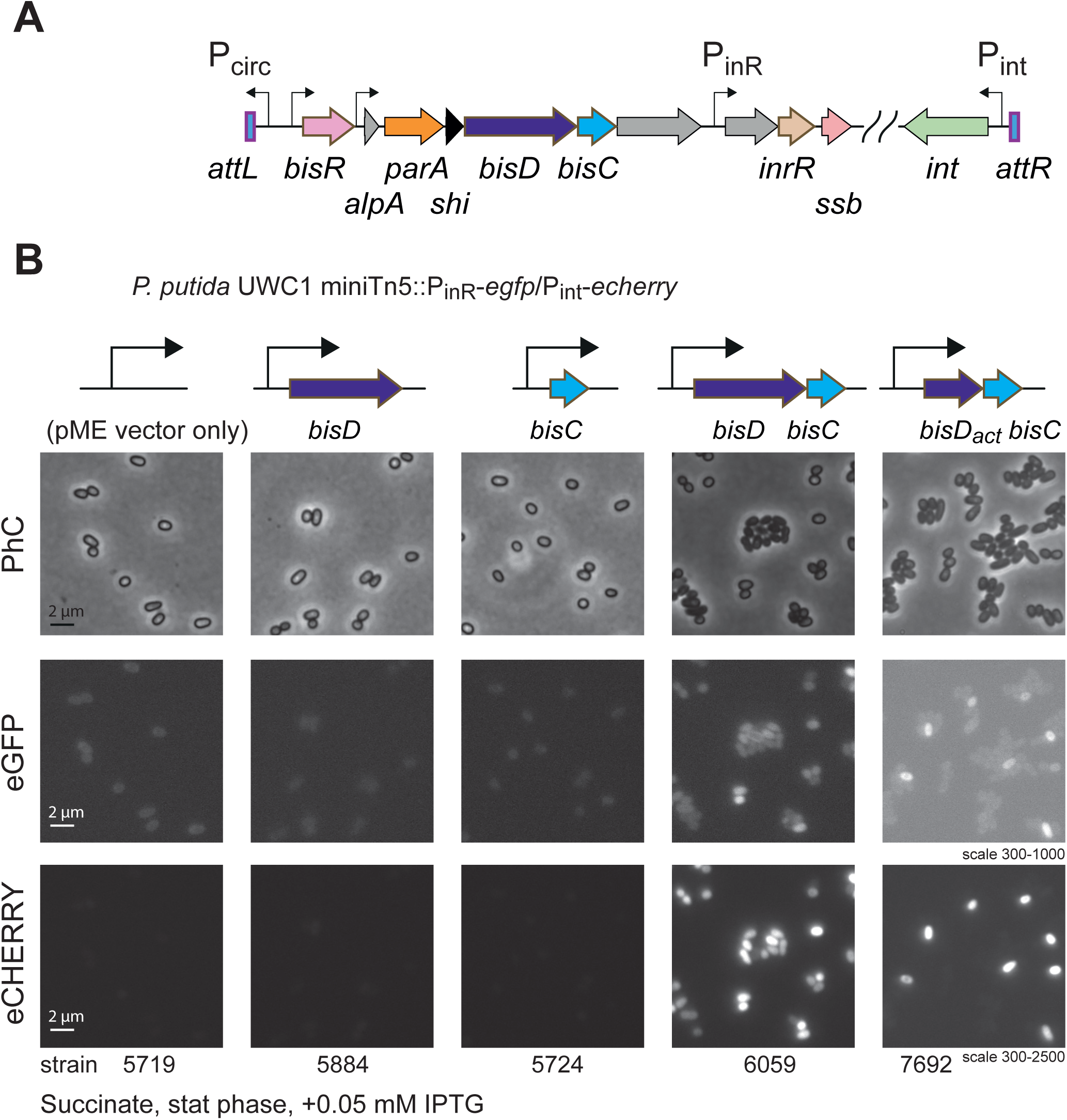
BisD-activator domain in conjunction with BisC is sufficient to activate expression from late ICEclc transfer competence genes. (A) Overview of the *bisD-bisC* locus on the integrated ICE*clc* in orientation to the *attL* and *attR* sites. Relevant gene names used in the text are indicated below the colored arrows representing their open reading frames and orientation. Hooked arrows point to previously identified ICE-promoters; those upstream of *bisR*, *alpA*, *inrR* and *intB13* are only activated in transfer competent cells. (B) Micrographs of *P. putida* without ICE*clc* but with a single copy integrated fluorescent reporter for activation of the *inrR*- and *int*-promoters, and transformed with pME6032-plasmid constructs containing either *bisD*, *bisC*, both *bisD-C* or only *bisD*-activator domain part with *bisC*, under control of the P_tac_-promoter. Cells were grown on succinate to stationary phase to allow coactivation of ICE-promoters by RpoS, and induced with IPTG to allow expression from P_tac_. Micrographs show phase-contrast (PhC), eGFP fluorescence (for P_inR_-expression), or eCherry fluorescence (for P_int_-expression). Relevant strain numbers for Table S1 are indicated. Fluorescence images scaled to the same minimum-maximum intensity per row. Note the subpopulation-dependent expression for strains with induced *bisD_act_-bisC*, but not with *bisDC*.

### BisD binds ICEclc DNA as distinct foci

The clamp-like predicted structure of the BisD dimer (Fig. 1) would support BisD-accumulation on specific DNA target sites similar to ParB/*parS*. To confirm this experimentally, we fused BisD and BisC individually with a fluorescent tag by allelic replacement on ICE*clc* and studied their subcellular localization in *P. putida* by epifluorescence microscopy. An N-terminal fusion of mCherry to BisD (separated by a short linker), and a carboxy-terminal fusion of the reading frame of *bisC* to *mCherry* (and a linker) were constructed in *P. putida* carrying a single ICE*clc* copy (Fig. 3A, B). Strains with either translational reporter fusion showed indistinguishable ICE*clc* self-transfer frequencies to wild-type ICE*clc*, indicating that the fusions did not impair their function in the ICE activation and conjugation process (Fig. 3C). Inducing expression of *mCherry-bisD* in *P. putida* ICE*clc* with help of an overexpressed ectopic copy of *bisR* on a plasmid (Carraro, Richard, et al., 2020) resulted in individual cells displaying between 1–4 distinct fluorescent foci (Fig. 3A). Cells with 1–2 foci likely represent mCherry-BisD proteins binding to the integrated ICE, but those with more than 2 foci are indicative for binding to excised ICE*clc* DNA, which appears in stationary phase cells (Delavat et al., 2019). In contrast, expression of *bisC-mCherry* under the same conditions did not lead to visible fluorescent foci in regular epifluorescence microscopy, although very weak and diffuse foci were observed in confocal scanning high resolution microscopy (Fig. 3B, *Airyscan*). The latter may be the consequence of binding of BisC-mCherry or BisD-BisC-mCherry complexes to their respective promoters on chromosomal or excised copies of ICE*clc* in the cell. To study the requirements for BisD focus formation, we (over-) expressed the BisD translational fusion construct in various configurations in *P. putida* with or without ICE*clc* (Fig. 3D). As expected, inducing *mCherry-bisD* together with *bisC* from a plasmid in *P. putida* with the ICE, again yielded a small number of distinct fluorescent foci in individual cells in stationary phase (Fig. 3D). In absence of ICE*clc*, however, ectopic expression of *mCherry-bisD* and *bisC* led to homogenously fluorescent cells (Fig. 3D), indicating that foci formation is specific for ICE*clc* DNA. Induction of a single copy ectopically integrated *mCherry-bisD* in presence of the ICE, but without co-expression of *bisC*, resulted in individual cells displaying mostly a single focus – and again homogenous fluorescent cells in absence of the ICE (Fig. 3E). The reason for a single instead of multiple fluorescent foci as in Fig. 3D is due to the fact that BisD alone is insufficient to activate ICE excision and replication. Hence, only mCherry-BisD binding to the integrated ICE*clc* can be detected. Under conditions of its natural activation in stationary phase cells grown on 3-CBA, the proportion of cells displaying fluorescent foci is much lower, since limited to the 3-5 % of formed transfer competent cells (Sulser et al., 2022), but those again displayed between 1–3 foci (Fig. S2). This proportion is similar to estimates of the number of excised ICE copies in individual cells (Delavat et al., 2016) (see below) and suggests that each BisD focus represents an individual copy of chromosomal or excised ICE*clc*.

**Figure 3.**
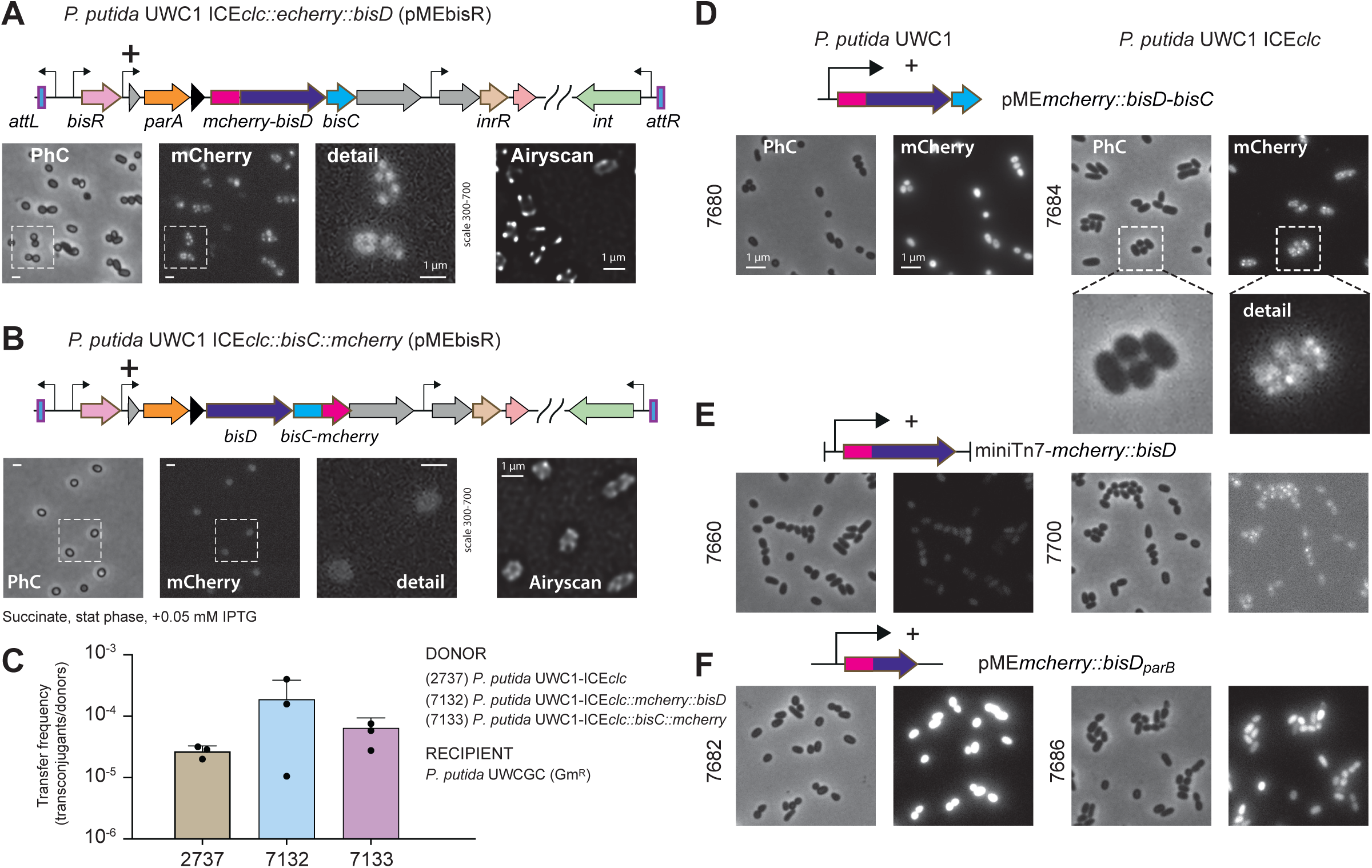
Fluorescent fusion proteins to BisD form ICE-DNA dependent foci in P. putida cells. (A) *P. putida* with integrated ICE*clc* on which the *bisD* reading frame is replaced by an in-frame *mcherry-bisD* translational fusion. Expression of the *alpA* promoter in all cells is induced by activating BisR from a transformed plasmid (pMEbisR). Micrograph annotation as in Fig. 2. Detail, magnification of the dotted region in mCherry fluorescence. Airyscan, independent imaged area showing individual cells with multiple fluorescent foci. Cells taken from stationary phase grown cultures on succinate, induced with IPTG. (B) *P. putida* with integrated ICE*clc* on which the *bisC* reading frame is replaced by an in-frame *bisC-mcherry* translational fusion, induced in all cells by overexpression from *bisR*. Same detail as in panel B, but without crisp fluorescent foci. (C) ICE*clc* self-transfer frequencies with *P. putida* wild-type ICE*clc*, or both translational fusions, as donors, showing no statistically significant differences. Hence, both translational fusion constructs are functional. (D) Ectopic expression of an mCherry-translational fusion to BisD in conjunction with BisC is sufficient to produce fluorescent foci in *P. putida* stationary phase cells induced with IPTG, but only in presence of the ICE. Micrograph details as in panel A. (E) Same as (D) but for a single copy integrated *mcherry-bisD* fusion. (F) Same as (D) and (E) but for ectopically expressed *mcherry* fused to a *bisD* with only the ParB domain, producing no fluorescent foci.

Finally, overexpressing BisD_par_, fused at its N-terminal end with mCherry, yielded no foci but homogenously bright fluorescence both in presence or absence of integrated ICE*clc* (Fig. 3F). The ParB-domain on BisD is thus not sufficient for foci formation, but requires the BisD_act_ domain for efficient DNA clamp formation.

### BisD recognizes two parS-like bisS sequences on ICEclc

As the epifluorescence microscopy experiments suggested specific recognition of ICE*clc* DNA by BisD, we attempted to map the *cis*-regulatory sequences on ICE*clc* required for BisD focus formation. We assumed that BisD_par_ may recognize specific DNA sequences similar to ParB binding to the cognate 16-bp palindromic *parS* sequences. *Pseudomonas* strains typically have 4 chromosomal *parS* sites with a consensus sequence of TGTTCCACGTGGAACN, at which the chromosomally encoded ParB accumulates (Fig. 4A). Screening ICE*clc* for the occurrence of (near)-palindromic *parS*-like sequences in the main ICE*clc* regulatory region, from *attL* to *ssb* (Fig. 4A), resulted in two candidate *bisS* sequences: CTTCGGAtTCCGAAG and GAGCTATCCCCnGGGGATAGCTC. The former (candidate *bisS-3*) comprising a 7-bp inverted repeat is located in a gene annotated as *orf96323* (Fig. 4A). The latter comprising an 11-bp inverted repeat is present in two instances on ICE*clc*, in the *inrR* gene and the *ssb* gene (Fig. 4A, named *bisS-1* and *bisS-2*, respectively). To test for the recognition of these putative target sites by mCherry-BisD, we cloned and inserted them in single copy using mini-Tn*5* integration on the chromosome of a *P. putida* strain lacking ICE*clc*. Induction of an ectopic copy of the *mCherry-bisD* gene from the *tac* promoter indeed resulted in (mostly) a single fluorescent focus appearing in stationary phase cells, but only when the insert included the *bisS-1* sequence or the *bisS-2* sequence (the reverse complement of *bisS-1*) (Fig. 4B). Cells with *bisS-3* did not produce fluorescent foci. These experiments indicated, therefore, that BisD specifically recognizes two *bisS* GAGCTATCCCCnGGGGATAGCTC sites on ICE*clc* that lay 2 kb apart.

**Figure 4.**
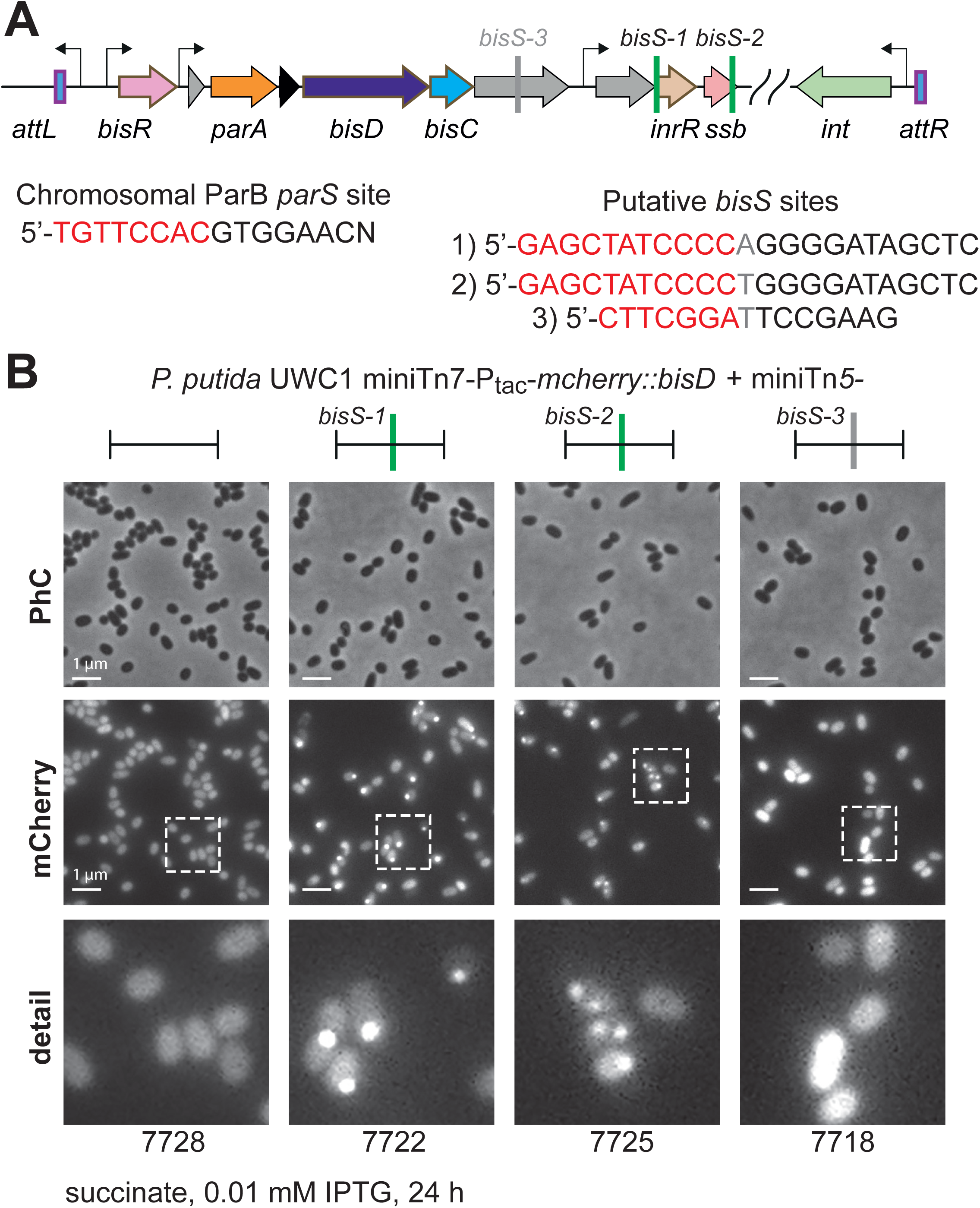
Fluorescent foci formation by BisD binding occurs at either of two regions on ICE*clc*. (A) Relevant gene region on ICE*clc* as before, but with three potential palindromic repeat BisD binding sites indicated. As a reference, the consensus *parS* site for the chromosomal ParB binding. (B) *P. putida* without ICE*clc* but with the single copy IPTG-inducible *mcherry-bisD* translational fusion and a single copy insertion of either empty mini-Tn*5* transposon, a cloned *bisS*-1, *bisS*-2 or *bisS*-3 fragment. Cells imaged after growth on succinate to stationary phase, in presence of IPTG to allow expression of mCherry-BisD. Note how strains with either *bisS*-1 or *bisS*-2 integrations show one or two fluorescent foci (as a result of chromosome replication), whereas those with *bisS*-3 show homogenous fluorescence. Images scaled to same minimum-maximum grey level per row. Detail shows the dotted mCherry zones. Relevant strain numbers indicated below the micrographs.

### BisDC localization and distribution on ICEclc in Pseudomonas putida

To investigate the distribution of BisDC on the ICE*clc* element, we employed a chromatin immunoprecipitation (ChIP) assay coupled to DNA sequencing (ChIP-seq) or quantitative PCR (qPCR). Expression of *bisD* and *bisC* alleles fused at their endogenous locus with an *mCherry* open reading frame enabled us to enrich the proteins by immunoprecipitation using anti-mCherry antibodies. *P. putida* ICE*clc* cells were grown until they reached stationary phase and then ICE*clc* transfer competence was artificially induced by overexpressing an ectopic copy of BisR, the transcription activator for *bisDC* (Carraro, Richard, et al., 2020). This artificial induction was used to increase the population of cells expressing BisDC, as natural activation upon growth with 3-CBA occurs in only 3-5 % of cells (Sulser et al., 2022). DNA immunoprecipitated with anti-mCherry serum was extracted, sequenced, and then mapped onto the *P. putida* ICE*clc* genome.

Analysis of the distribution pattern revealed a maximal enrichment of mCherry-BisD around the two copies of the *bisS* site on ICE*clc*, confirming our identification as BisD target sites (Fig. 5A). Strikingly, rather than showing local enrichment, the distribution of BisD was very broad, covering about half of the entire ICE*clc* but not the host chromosome. This broad coverage is similar to what is known from ParB at the chromosomal *parS* sites, albeit clearly more extensive, suggesting a shared mechanism of DNA association. The BisD distribution was asymmetric, extending unidirectionally towards the *attR* side of the ICE (Fig. 5B) and much less so into the chromosome flanking the ICE on the other side (i.e., beyond *attL*, Fig. 5B). The apparent bias of BisD spreading towards the centre of ICE*clc* and away from the nearest chromosome flank, is likely not due to excision of ICE*clc* from the chromosome and subsequent replication as extrachromosomal element, since in one biological replicate no higher ratio of ICE*clc* reads compared to other parts of the genome was observed (indicating the absence of excision despite BisD association in this sample) (Fig. S3). The preference for spreading away to the ICE*clc* centre may be due to the direction of ICE core gene transcription (Fig. 5A), with BisD sliding following the transcription complex but hampering in the other direction. In contrast to BisD, BisC mapping showed much narrower peaks of enrichment with similar peak heights, which mostly localize to previously identified promoter regions of transfer competence genes under BisDC control (Sulser et al., 2022). Combined BisD/BisC-mapping also suggests there may be two further bistable promoters, one upstream of *iceB7* and the other upstream of the gene *orf84835.* Both regions had been noticed for absence of RNA readthrough (Gaillard et al., 2010), but not been verified with single copy promoter-reporter studies (Sulser et al., 2022). Notably, BisD showed increased occupancy near the BisC bound sites, indicating formation of BisDC complexes (analogous to FlhDC). We also quantified the enrichment of mCherry-BisD and BisC-mCherry by qPCR specifically for three ICE*clc* loci *bisS-1*, orf*96323* and *clcA* (Fig. S4). The results showed similar high enrichment of mCherry-BisD and BisC-mCherry on the *bisS-1* locus, and a lower enrichment on the downstream *96323* and *clcA* genes (1 kb and 80 kb, respectively, away from *bisS*) (Fig. S4), further corroborating the findings of the ChIP-seq. Of note, the overall enrichment of ectopically expressed mCherry-BisD was lower than that of the endogenously expressed, likely due to the absence of the positive feedback loop mediated by BisDC driving expression from its own promoter at the endogenous location (Fig. 5A). Notably, in this sample, the distribution of mCherry-BisD across *bisS* is more symmetrical, supporting the notion that the direction of gene transcription leads to the asymmetry in BisD spreading at advanced stages of ICE*clc* activation (Fig. 5A). Altogether, the results suggest a BisD-DNA association similar to ParB proteins at *parS* sites: local loading at *bisS* sites and asymmetric spreading to adjacent DNA over long genomic distances (50 kb and more).

**Figure 5:**
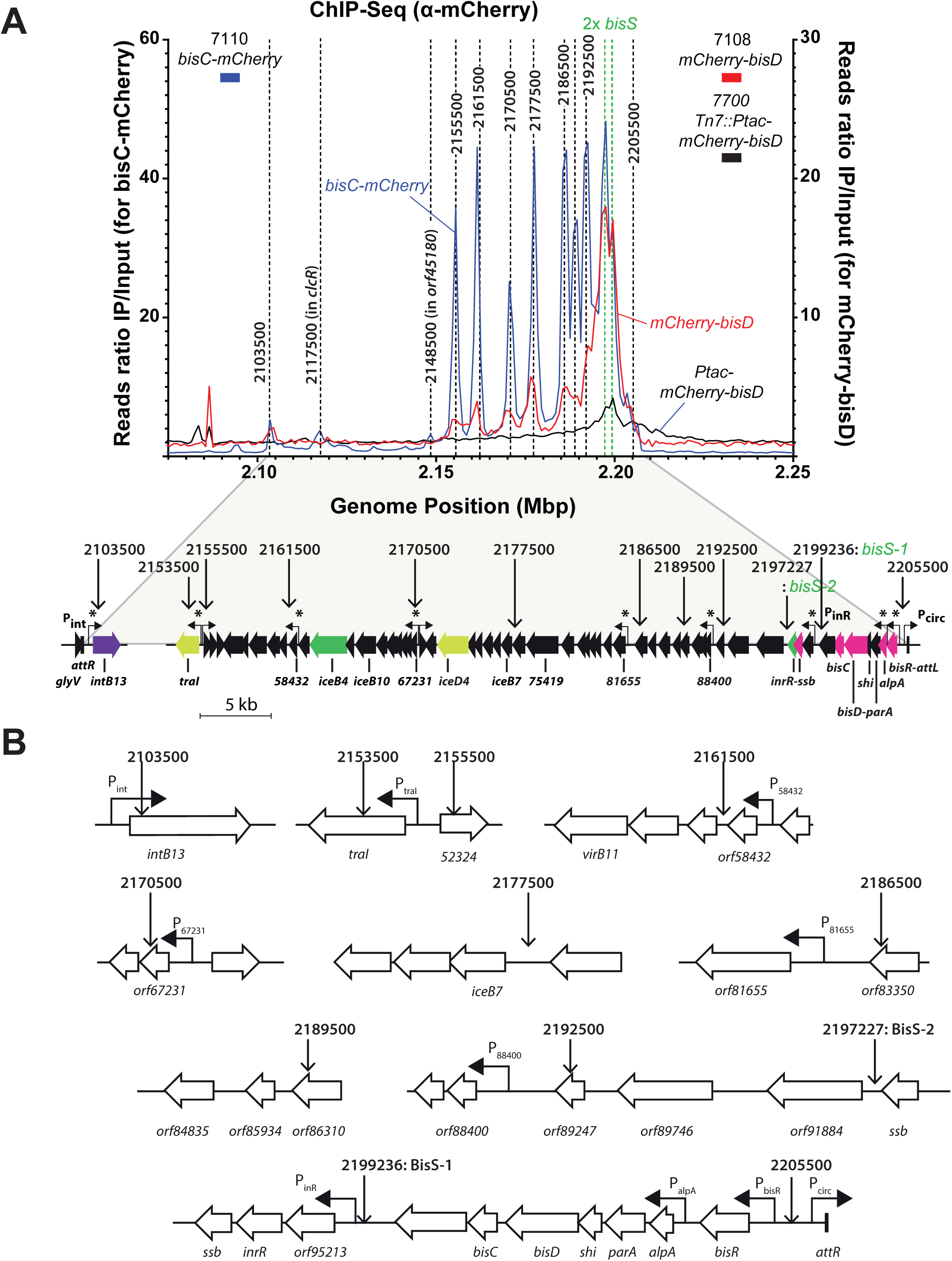
Identification of BisD- and BisC-binding sites on ICE*clc* in *P. putida*. (A) Chromatin-immunoprecipitation coupled to deep sequencing (ChIP-Seq) using α-mCherry antibody. The panel shows an overlay of the BisD- and BisC-mCherry distribution profiles (each IP sample DNA is normalized to its respective input DNA) in a zoomed-in region of the integrated ICE*clc* including the two identified *bisS* sites. Relevant ICE*clc* genes and their orientations are shown by colored arrows with names below. Hooked arrows with a star correspond to previously identified transfer competence promoters by Sulser et al. (Sulser et al., 2022). The full distribution profile on the entire *Pseudomonas putida* genome is shown in Fig. S2. All ChIP-Seq profiles were split into 1 kb bins. (B) Detail of identified peak regions (number gives deduced chromosome position) in relation to location and orientation of relevant ICE*clc* genes (as open arrows - gene names or orf numbers indicated). Hooked arrows point to previously identified transfer competence promoters expressed in the same 3-5% subset of cells in stationary phase.

### BisD binds NTPs via the conserved GxxR motif with varying affinities

Given the notable similarities in structure and chromosomal distribution between BisD and ParB proteins, we investigated whether BisD binds any ribonucleotides, similarly to ParB proteins binding to CTP. Knowing that ParB is particularly important during exponential growth, while ICE*clc* BisDC is activated only in stationary phase cells, we wondered whether ParB and BisD could utilize and sense different nucleotide cofactors. The AF3-predicted structure of BisD_par_ suggests the presence of a pocket that may potentially accommodate a ligand. To explore this, we employed isothermal titration calorimetry (ITC) to quantify the binding interactions between BisD and the four standard ribonucleotides: CTP, ATP, GTP, and UTP. BisD was recombinantly expressed, purified, and subjected to gel filtration immediately prior to ITC analysis. Strikingly, the experimental results revealed a binding affinity of BisD in the range of ≈ 2 μM (Kd) for CTP and 13 μM for ATP, while demonstrating somewhat lower estimates for the affinities for UTP and GTP (Kd ≈ 21 μM and ≈ 64 μM, respectively) (Fig. 6). CTP and ATP binding was abolished in the BisD (R95A) mutant, indicating that this residue in the GxxR motif contributes to the nucleotide binding pocket (Fig. S5).

**Figure 6:**
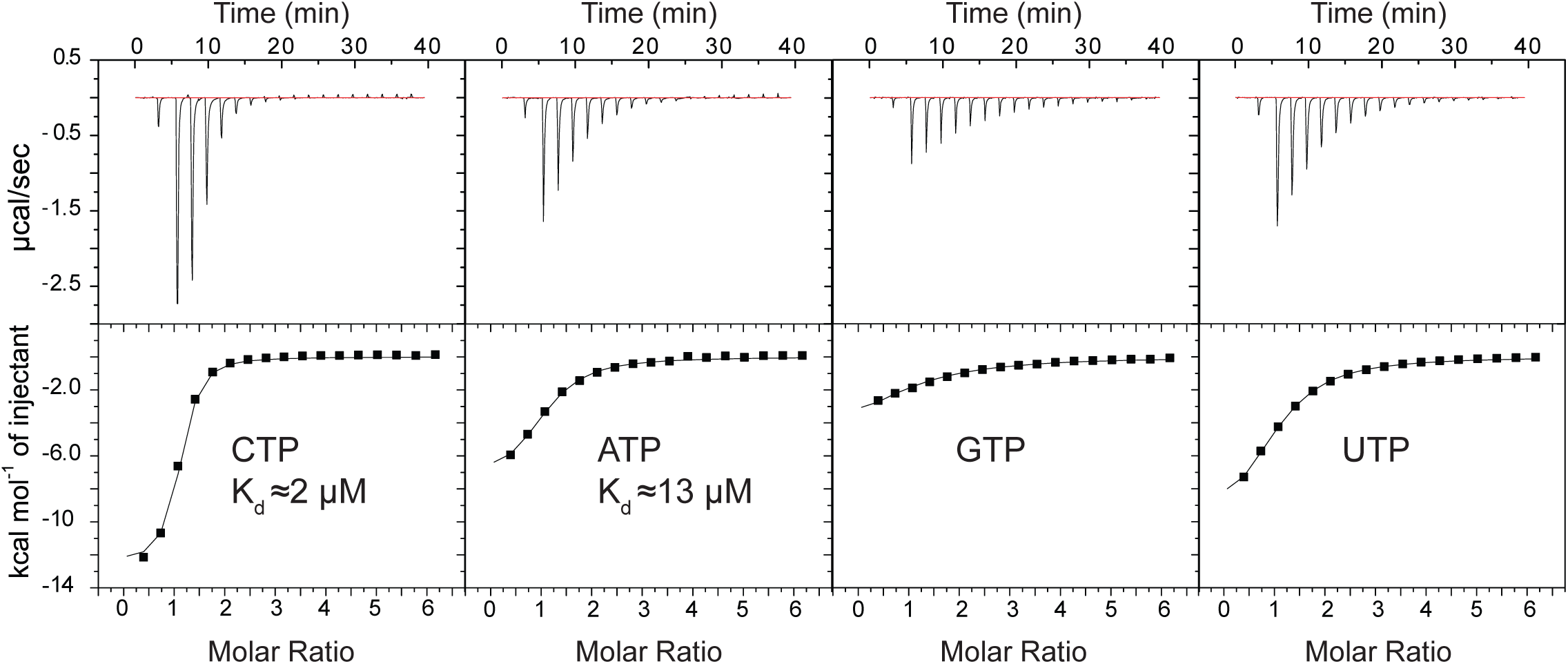
BisD-NTP affinity measurements by isothermal titration calorimetry (ITC). Wild-type BisD, at a monomer concentration of 60 μM, was injected with 2 mM NTP solution. The Kd from a typical experiment is given. See Materials and Methods for the experimental details.

### BisD shows appreciable levels of CTP and ATP hydrolysis

We then checked if BisD can hydrolyze any of the ribonucleotides it binds to. We utilized endpoint colorimetric detection of free inorganic phosphate released by nucleotide hydrolysis in the Malachite green assay. During one hour of incubation at 25°C, BisD showed low but appreciable levels of hydrolysis of CTP but not of ATP, GTP, or UTP (Fig. 7). CTP hydrolysis was slightly stimulated upon addition of an ICE*clc* 40-bp DNA fragment containing a *bisS-1* site but not with an unrelated DNA sequence of the same length. This result suggests that BisD, although potentially binding all ribonucleotides, selectively hydrolyses CTP, a reaction which is stimulated by *bisS* DNA.

**Figure 7:**
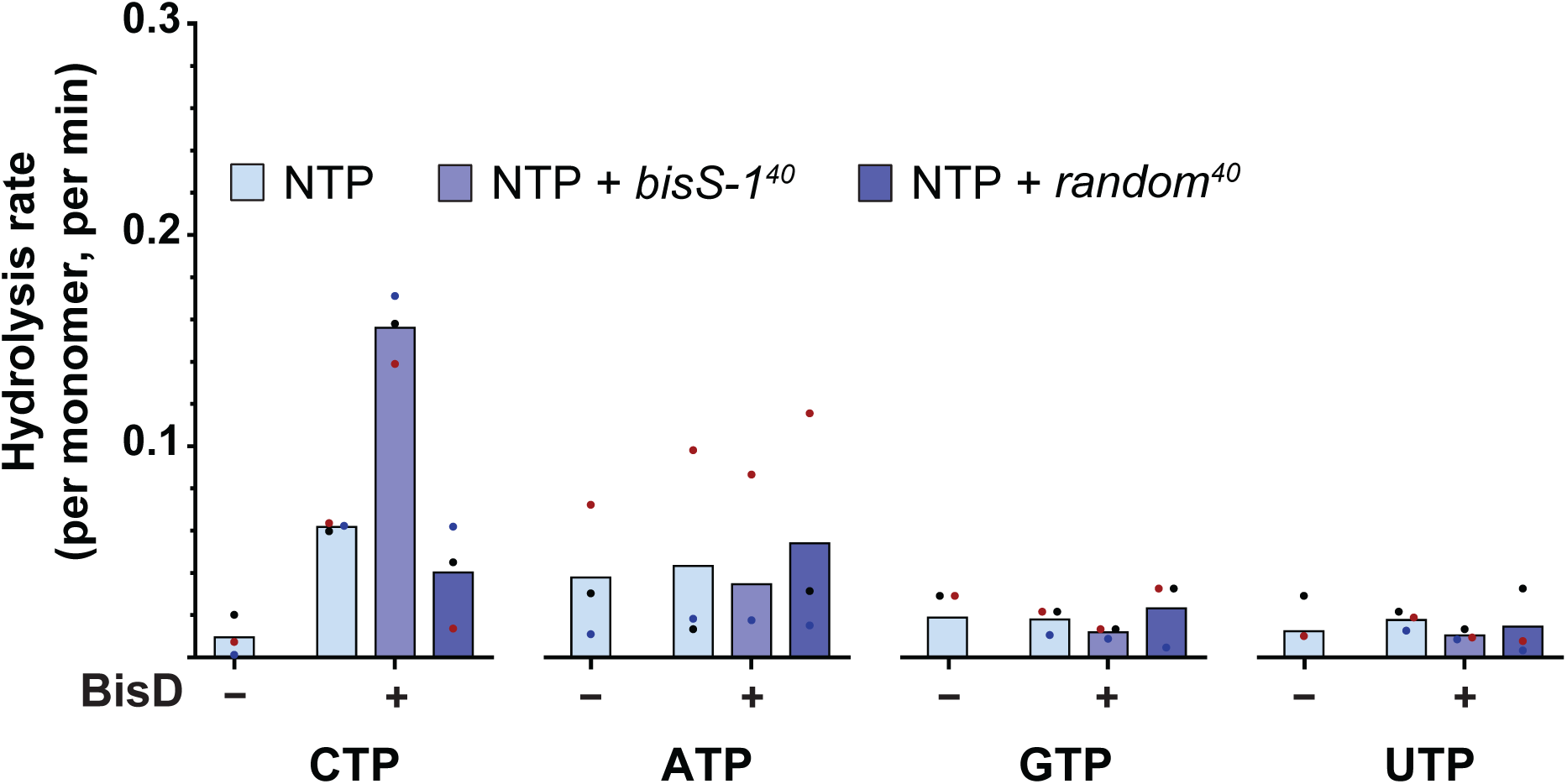
NTP hydrolysis rate by BisD. Measured by colorimetric detection of inorganic phosphate using Malachite Green Assay. 10 μM of protein was incubated with 1 mM NTP with or without 1 μM DNA_40_. Mean values are calculated, and individual data points are shown as dots

## Discussion

We characterize here for the first time a unique natural hybrid protein formed from a domain with ParB-function and a second domain with a transcription regulatory function. This protein, named BisD, occurs on the ICE*clc* element in *P. putida* and *P. knackmussii*, where together with BisC, it was shown to be involved in activation of ICE*clc* transfer competence (Carraro, Richard, et al., 2020). The N-terminal ParB-like domain of BisD had been noticed previously (Reinhard et al., 2013), but we firmly demonstrate here the hallmarks of known ParB-function: (i) predicted sequence and structural similarity to ParB proteins with conserved residues forming the CTP binding pocket, (ii) recognition of and spreading from specific DNA sites (*bisS*), and (iii) nucleotide hydrolysis in presence of *bisS*-DNA. We also show that BisD has a gene activation function, which is located in its C-terminal domain. The BisD activator domain can function independently of its ParB-domain (when overexpressed) but needs BisC for activation of ICE*clc* transfer competence promoters. BisD and BisC enrich and colocalize to promoter regions on ICE*clc* that have been shown to bistably express exclusively in transfer competent *P. putida* cells (Sulser et al., 2022), and fluorescently tagged BisD forms distinct foci that are dependent on the presence of a *bisS* site. The BisD/BisC-gene pair occurs widely on ICEs of the same family as ICE*clc*, suggesting the dual function to have selective benefits for ICE-transfer.

Our first hint for enrichment of BisD on ICE-DNA came from microscopy images of *P. putida* with a single integrated ICE*clc* grown to stationary phase on 3-CBA. Under these conditions, the highest proportion of transfer competent cells is formed, and only those cells displayed clear mCherry-BisD fluorescent foci. Activation of transfer competence can be achieved more widely by inducing expression of the BisR-activator in stationary phase cells, in which case all individual cells displayed fluorescent foci. Between one and up to 4 foci could be distinguished, which is suggestive for mCherry-BisD enriching on a single integrated (one focus) and on excised ICE*clc* copies, of which we previously showed that multiple can be replicated in a single stationary phase cell (Delavat et al., 2019). By ectopic and heterologous expression in absence of the ICE, we demonstrate that this binding is dependent on one of two specific sites on the ICE (named *bisS1* and *bisS2*) and only requires BisD itself. In contrast to BisD, BisC-mCherry did not form noticeable foci, indicating that foci visibility must be due to larger quantities of mCherry-BisD protein accumulating on the DNA. The BisD-ParB domain alone is not sufficient to form foci.

Binding of BisD was confirmed by ChIP-seq profiling, which showed BisD enrichment at the two *bisS* loci but also further broad local enrichments on ICE*clc* DNA that includes all known bistable promoter regions activated by BisDC in transfer competent cells. Inter alia: ChIP profiling suggested two further BisDC-activated promoters that we had not detected in a previous bistable promoter screening (Sulser et al., 2022), but which may thus constitute *bona fide* transfer competent promoters. The broad mCherry-BisD enrichment peaks colocalized precisely with those detected by ChlP-Seq with BisC-mCherry, which were sharper and more pronounced, possibly because their binding site is more defined. Gene deletions showed that the C-terminal domain of BisD (BisD_act_) is sufficient for transcription regulation of the ICE*clc* promoters (when overexpressed). Our results, therefore, indicate a model where BisD first loads at *bisS* sites and would then spread as a ParB-like clamped protein to adjacent DNA. Promoter regions are then recognized either because BisD helps to recruit BisC and find or enhances BisC-DNA specificity, or BisD spreading and scanning the adjacent DNA until it finds an already promoter-associated BisC protein, after which transcription activation occurs by BisD_act_ interactions to BisC (Fig. 8). This is reminiscent of the ParB protein KorB from *E. coli* plasmid RK2, which represses genes together with the KorA partner protein. KorAB regulation, however, acts over significantly shorter distances (several hundred base pairs) between the KorB loading sites and the respective promoters (McLean et al., 2024).

**Fig. 8.**
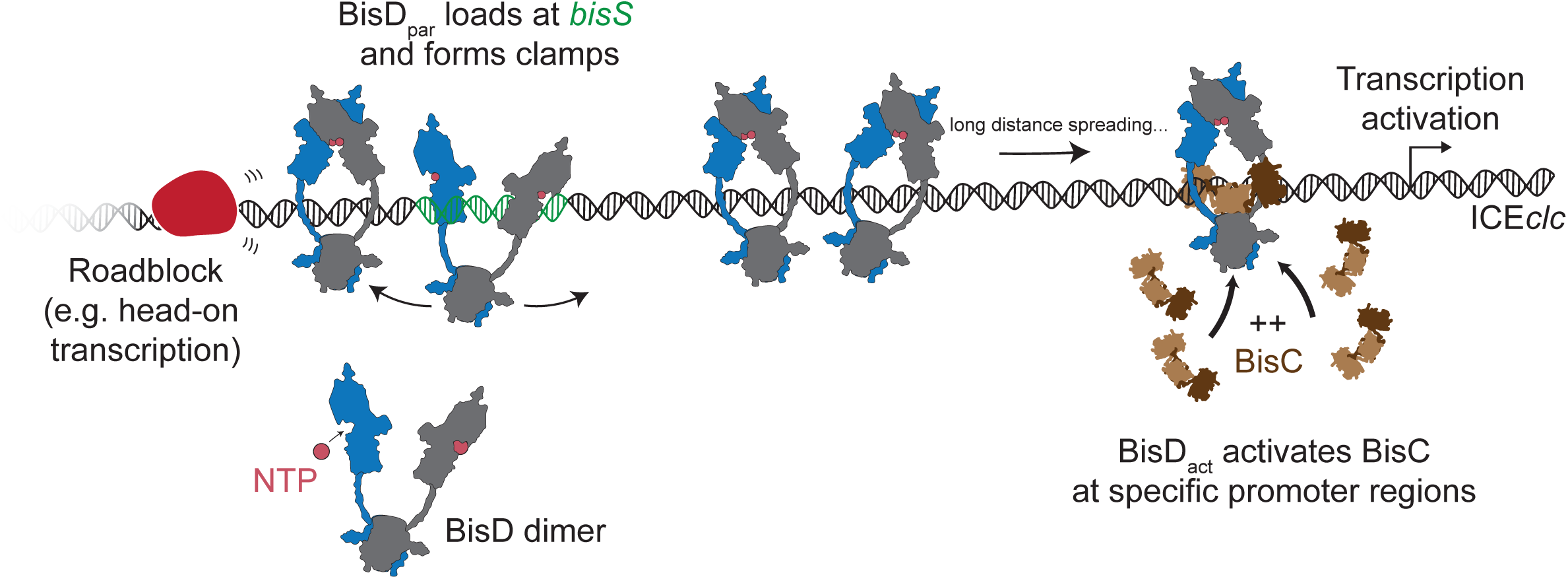

Purified His_6_-BisD indeed binds any of the four nucleotides, with CTP having highest affinity in a similar range as shown for ParB proteins of various origins. We showed that a mutation in the conserved nucleotide binding pocket GxxR and ENxxR in the BisD_par_ domain abolishes nucleotide binding. We could also show that purified His_6_-BisD hydrolyses CTP, which is stimulated upon the presence of 40-bp double-stranded *bisS* DNA fragments. These are similar characteristics as have been observed for purified ParB proteins and their *parS* site. In the current model *parS* catalyses a conformational change of ParB dimers into clamps that slide around the DNA, and this forms the active CTPase. By analogy it would thus be conceivable that interaction of BisD with its *bisS* leads to BisD-dimer clamps that entrap the DNA and spread to adjacent regions. Curiously, BisD spreading seems rather asymmetric and proceeds primarily ‘into’ the ICE following the transcription direction of most of the ICE*clc* core genes, and not across the ICE *att* junctions (Fig. 5). This asymmetry is less dramatic when BisD is ectopically expressed, implying that it arises at least partly during ICE*clc* activation, possibly by transcription pushing BisD clamps in one direction and/or preventing sliding in the other direction. Considering the large distance of several BisD-targeted promoters from the BisD loading sites (*bisS*), it is tempting to speculate that spreading of BisD from *bisS* to the target promoter establishes a temporal order of gene activation with more distant genes being activated later. Unfortunately, the poor synchrony of ICE*clc* activation across a population of cells (Sulser et al., 2022) does not allow us to test this hypothesis directly, but the decay in BisD binding (i.e., the lower proportion of BisD-bound reads) from position 2192500 towards the centre of the ICE (Fig. 5A) mirrors the decrease in relative ICE-transcript abundance previously measured in RNAseq (Sulser et al., 2022).

CTP hydrolysis would eventually lead to opening of the BisD-clamp and its dissociation from the DNA, as has been reported for canonical ParB proteins (Antar et al., 2021; Jalal et al., 2021; Osorio-Valeriano et al., 2021). The rate of CTP hydrolysis of BisD is slower than that of chromosomal ParB of *Bacillus subtilis* (around 8 to 10-fold lower), but other ParB-like proteins have been reported to exhibit a slow hydrolysis rate (Antar & Gruber, 2023; Jakob et al., 2024) or even lack hydrolysis activity entirely (Osorio-Valeriano et al., 2019). Possibly, the slow CTP hydrolysis rate of BisD leads to a more extensive coverage of ICE*clc* DNA, which is advantageous for its regulatory mechanism of coactivating some 10 downstream ICE promoters. The 50-kb coverage by BisD is more than has been measured for other (wild-type) ParB proteins and their respective chromosomal target DNA. Slow hydrolysis would come with a risk of accumulating off-target BisD clamps, but in case of the ICE*clc* this seems to be avoided, perhaps again because BisD expression is restricted to stationary phase transfer competent cells that do not undergo cell division.

The ParB-like mechanism of BisD of sliding across DNA from the *bisS* sites is reminiscent for a partitioning-like mechanism. Indeed, a gene encoding for a ParA-like protein exists upstream of *bisD* on the ICE (Fig. 2A). The question is then justified as to why ICE*clc* would need its own partitioning system, since previous results indicated that transfer competent cells are locked in a bistable state of ICE conjugation, and cannot go back to regular cell division because of replication fork stalling by ParA and the protein Shi (also encoded on ICE*clc*, Fig. 2A) (Reinhard et al., 2013; Takano et al., 2019). Either the BisD-mediated partitioning is then a relict to distribute excised and replicated ICE-copies among daughter cells, or somehow the ParA-BisD-*bisS* system acts to transport excised ICE*clc* DNA to the type IV secretion system conjugation complexes. Daughter cell partitioning of excised ICE-DNA is not very likely, as previous single cell microscopy data indicated that dividing transfer competent cells frequently lose the ICE (Delavat et al., 2016), and their cell division is inhibited anyway by the Shi-system (Takano et al., 2019). Active transport of excised ICE*clc* DNA molecules to conjugation complexes could be advantageous for transfer and avoid clogging of multiple ICE molecules with multiple conjugation pores around the cell envelope (Daveri et al., 2023), but this would require some sort of recognition of the ICE-DNA-BisD complex by the type IV secretion system. In this context it is interesting to note that plasmid-expressed ParA-mCherry in *P. putida* (without ICE) locates to the nucleoid and concomitantly expressed Shi-eGFP localizes as foci in the cell envelope (Takano et al., 2019). In addition to their inhibition of chromosome replication, ParA and Shi might thus be involved in pushing ICE-DNA away from the nucleoid to conjugation pores, of which there are multiple in transfer competent cells (Daveri et al., 2023).

ParA-BisD-BisC may thus be the result of an evolutionary fusion of two functionally unrelated systems resulting in a bifunctional module involved in (i) the coordinated activation of target bistable ICE promoters ensuring formation of the conjugation transfer system exclusively in transfer competent cells, and (ii) active partitioning together of excised ICE. In this context, the fusion of functionally distinct domains in BisD confers an advantage by temporally coupling the activation of gene expression with the transfer of ICE*clc* to other cells. The dual function of BisD could ensure a coordinated progression from the development of a transfer-competence state to the actual process of transferring ICE*clc*. Future work is needed to approach these hypotheses.

## Acknowledgements

We are grateful to members of the van der Meer and Gruber laboratories for stimulating discussions, technical advice and help, and critical comments on the manuscript. We thank Laura Chamera Rendueles for her help in preliminary *bisDC* cloning and analysis.

## Author Contributions

H.A., protein purification and biochemical experiments. H.A., N.C., chromatin immunoprecipitation. N.C., *Pseudomonas* strain construction, culturing, and imaging. All authors prepared and revised the manuscript text and figures.

## Data and Materials Availability

All data needed to evaluate the conclusions in the paper are present in the paper and/or the Supplementary Materials. Raw sequencing data obtained in this work will be deposited to NCBI-SRA.

## Materials and methods

### Bacterial strains and culturing

Bacterial strains and plasmid constructions used in this study are shortly described in Table S1. Strains were routinely grown in Luria broth (10 g l^−1^ Tryptone, 10 g l^−1^ NaCl and 5 g l^−1^ Yeast extract, LB Miller, Sigma Aldrich) at 30 °C for *P. putida* and 37 °C for *Escherichia coli* in an orbital shaker incubator and were preserved at –80 °C in LB broth containing 15 % (v/v) glycerol. Reporter assays and transfer experiments were carried out with cells grown in type 21C minimal media (Gerhardt et al., 1981) supplemented with 10 mM sodium succinate or 5 mM 3-chlorobenzoate (3-CBA). Antibiotics were used at the following concentrations: ampicillin (Ap), 100 µg ml^−1^ for *E. coli* and 500 µg ml^−1^ for *P. putida*; gentamycin (Gm), 10 µg ml^−1^ for *E. coli*, 20 µg ml^−1^ for *P. putida*; kanamycin (Kn), 50 µg ml^−1^; tetracycline (Tc), 12 µg ml^−1^ for *E. coli*, 100 µg ml^−1^ or 12.5 µg ml^−1^ for *P. putida* grown in LB or MM, respectively. Transcription was induced from *P_tac_* by supplementing cultures with 0.05 mM isopropyl β-D-1-thiogalactopyranoside (IPTG; or else at the indicated concentrations).

### Molecular biology

Relevant DNA fragments from ICE*clc* for subsequent cloning or ectopic expression were obtained by PCR, with primers described in Table S2. PCR products were purified using Nucleospin Gel and PCR Clean-up kits (Macherey-Nagel) according to manufacturer’s instructions. PCR-amplified DNA for cloning was tailored either by specific restriction enzyme digestion (all restriction enzymes purchased from New England Biolabs) and ligated to the corresponding vector by overlap using the Vazyme ClonExpress II one-step method. As vectors we used the mini-Tn*5* (Martinez-Garcia et al., 2011) or mini-Tn*7* delivery plasmids (Choi et al., 2005), or the *lacI^q^-*P controllable expression shuttle vector pME6032 (Heeb et al., 2000). Plasmids were first constructed in *E. coli*, purified from there using a Nucleospin Plasmid kit (Macherey-Nagel) according to manufacturer’s instructions, and then transformed into the appropriate *P. putida* UWC1 or UWC1-ICE*clc* hosts by electroporation as described by (Dower et al., 1988) in a Bio-Rad GenePulser Xcell apparatus set at 25 µF, 200 V and 2.5 kV for *E. coli* and 2.2 kV for *P. putida* using 2-mm gap electroporation cuvettes (Cellprojects). All constructs were verified by DNA sequencing on purified plasmid-DNA (Eurofins). Mini-transposed DNA insertions in the *P. putida* genome were verified by PCR and by appropriate marker selection.

### Construction of mCherry-BisD and BisC-mCherry fusions

Translational fusions were constructed between mCherry and BisD, and between BisC and mCherry, by allelic replacement on a single ICE*clc* copy integrated in the chromosome of *P. putida* UWC1. First we amplified the *mcherry* gene without stop codon and with a 45-bp extension for a linker peptide, as described in (Miyazaki et al., 2012), which was fused downstream to a ca. 700-bp amplified 5’-gene fragment of *bisD* and upstream to a ca. 700-bp fragment covering the *shi* and part of *parA-*gene, such that the start codon of *mcherry* becomes the native start of *bisD.* Similarly, a DNA fragment was constructed carrying *bisC* without its stop codon, followed by the linker and *mcherry* open reading frame, and a ca. 700-bp recombination region directly downstream of *bisC*. Both assembled fragments were cloned individually into plasmid pJP5903, introduced into *P. putida* UWC1-ICE*clc*, and selected for single recombination events. Purified single recombinants were then induced by transformation with plasmid pSW and 3-methylbenzoate to force unique chromosome breaks and facilitate recovery of seamless in-frame double recombinants, as described by (Martinez-Garcia et al., 2011). Double recombinants were purified, cured from pSW and verified by PCR for the correctness of the intended construction. The same fusion fragments were then also cloned into pME6032 or into mini-Tn*5* or mini-Tn*7* vectors for ectopic screening.

### Conjugative transfer assays

ICE*clc* transfer was tested with 96-h-3CBA-grown donor and 24-h-succinate-grown recipient cultures. Cells were harvested by centrifugation of 1 ml (donor) and 2 ml culture (recipient, Gm-resistant *P. putida* UWCGC) for 3 min at 1200 × *g*, washed in 1 ml of MM without carbon substrate, centrifuged again and finally resuspended in 20 µl of MM. Donor or recipient alone, and a donor-recipient mixture were deposited on 0.2–µm cellulose acetate filters (Sartorius) placed on MM succinate agar plates, and incubated at 30 °C for 48 h. The cells were recovered from the filters in 1 ml of MM and serially diluted before plating. Cell suspensions were serially diluted (10-fold), and dropped and dried on MM agar plates containing appropriate antibiotics and/or carbon source (3-CBA). Transfer frequencies are reported as the mean of the exconjugant colony forming units compared to that of the donor in the same assay.

### Epifluorescence microscopy

Expression and localization of fluorescent reporter signals in *P. putida* cultures were quantified by epifluorescence microscopy on individual cells. Cells (150 µl aliquots) were sampled from IPTG-induced cultures or from stationary phase cultures on 3-CBA, which were briefly vortexed (30 s at max speed) to disrupt potential cell aggregates, and 5 µl drops were then spread on agarose-precoated glass slides (ca. 1 mm thick, using 1% agarose in 21C MM). Glass slides were covered with a regular 24×50×0.15 mm cover slip (Menzel-Gläser), and imaged in upright position with a Zeiss Axioplan II or in inverted position with Nikon Eclipse Ti-E microscope, as previously described (Carraro, Richard, et al., 2020; Daveri et al., 2023). Cells were imaged in phase contrast (PhC, 40 ms exposure), mCherry (500 ms) and eGFP fluorescence wavelengths (500 ms), and saved as 16-bit TIF-files. For better resolution, we used a ZEISS LSM 980 Airyscan 2 microscope, imaging cells in 14 subsequent Z-layers of 0.17 µm each in either GFP or e/mCherry channels (314 ms exposure), which were deconvoluted using the Zeiss system software with default settings. Fluorescence images for display were scaled to the same brightness in ImageJ (Schneider et al., 2012) as indicated, saved as 8-bit gray TIF-files, cropped to the display area and saved as 300 dpi resolution in Adobe Photoshop (Adobe vs. 2023).

### Chromatin immunoprecipitation

Relevant *P. putida* cultures were grown in 10 mL of type 21C minimal media with the addition of IPTG to 0.05 mM concentration at 30°C for 16 h overnight until they reached stationary phase. Subsequently, 1 mL of fixation solution F (comprising 50 mM Tris-HCl at pH 7.4, 100 mM NaCl, 0.5 mM EGTA at pH 8.0, 1 mM EDTA at pH 8.0, and 10% formaldehyde) was introduced and the mixture was left to incubate at room temperature for 30 minutes with occasional hand shaking. Following this, the cells were collected through centrifugation and washed using 1×PBS. The biomass of each specimen was equalized to represent 2 OD_600_ units (equivalent to 2 mL at OD600 = 1) and was then mixed with 1 mL of TSEMS lysis solution, containing 50 mM Tris/HCl pH 7.4, 50 mM NaCl, 10 mM EDTA (stock solution at pH 8.0), 0.5 M sucrose, protease inhibitor cocktail (Sigma), and 6 mg/mL lysozyme (Sigma). The cell suspension was incubated at 37 °C for 30 minutes with vigorous shaking. The resulting protoplasts were centrifuged, washed in 2 mL of TSEMS, centrifuged and resuspended in 1 mL of the same buffer, divided into three aliquots, again centrifuged to pellet the cells, which were flash frozen in liquid nitrogen, and stored at –80°C for later use.

One individual pellet from each sample was resuspended in 2 mL of buffer L, containing 50 mM HEPES-KOH at pH 7.5, 140 mM NaCl, 1 mM EDTA at pH 8.0, 1% Triton X-100, 0.1% SDS, 0.1 mg/mL RNase A, and protease inhibitor cocktail (Sigma). This mixture was then transferred to 5 mL round-bottom tubes. The cells were subjected to three cycles of sonication for 20 seconds each using a Bandelin Sonoplus with an MS72 tip (settings: 90% pulse and 35% power output). The lysates were moved to 2 mL tubes and centrifuged for 10 minutes at 21,000 × *g* at 4°C. From the supernatant, 800 µL was used for immunoprecipitation (IP), and 200 µL was set aside as the whole cell extract (WCE), which was then frozen at –20°C for future use.

For the IP process, anti-mCherry antibodies (Takara cat. # 632496) were initially mixed with Protein G attached dynabeads (Invitrogen) in a 1:1 ratio and incubated for 2 hours at 4°C on a rotating platform. The beads were then washed in buffer L, and 50 µL were added to each sample tube. The samples were incubated with the beads for 2 hours at 4°C with rotation. Subsequent to the incubation, the beads in the samples underwent a sequence of washes using buffer L, buffer L5 (buffer L with 500 mM NaCl), buffer W (containing 10 mM Tris-HCl at pH 8.0, 250 mM LiCl, 0.5% NP-40, 0.5% Na-Deoxycholate, and 1 mM EDTA at pH 8.0), and finally buffer TE (10 mM Tris-HCl at pH 8.0 and 1 mM EDTA at pH 8.0). Post-washing, the beads were resuspended in 520 µL of TES buffer (50 mM Tris-HCl at pH 8.0, 10 mM EDTA at pH 8.0, and 1% SDS). The previously frozen WCE was thawed, and 300 µL of TES and 20 µL of 10% SDS were added. Both the IP and WCE mixtures were incubated overnight at 65°C with vigorous shaking to reverse formaldehyde cross-links. The following day, DNA was purified from these samples using phenol-chloroform extraction. This involved vigorous mixing of the samples with 500 µL of phenol (buffered with 10 mM Tris-HCl at pH 8.0 and 1 mM EDTA), followed by centrifugation. Then, 450 µL of the upper phase was mixed with an equal volume of chloroform and centrifuged again. Finally, 400 µL of the aqueous layer was used for DNA precipitation using 1 mL of 100% ethanol, 40 µL of 3M sodium acetate, and 1.2 µL of GlycoBlue, incubated for 20 minutes at –20°C. The samples were then centrifuged, and the DNA pellets were dissolved and purified using QIAGEN PCR purification kit, eluted in 50 µL of EB buffer.

For qPCR analysis, 1:10 and 1:1000 dilutions of the IP and WCE DNA were prepared in water, respectively. Each 10 µL reaction, set up in duplicate, included 5 µL of Takyon SYBR MasterMix, 1 µL of a 3 µM primer pair (Table S2), and 4 µL of DNA. These were run in a Rotor-Gene Q machine (QIAGEN). Raw data were analyzed using the PCR Miner server (http://ewindup.info).

The sequencing of IP-DNA was conducted by the Lausanne Genomic Technologies Facility at the University of Lausanne. The DNA was initially fragmented using a Covaris S2 sonicator to sizes between 300–400 bp. DNA libraries were prepared using the Ovation Ultralow Library Systems V2 Kit (NuGEN), including 15 cycles of PCR amplification. Sequencing was performed on a NovaSeq (Illumina), yielding around 43 million pair-end sequence reads per sample. The reads were then processed by mapping them to *P. putida* UWC1-ICE*clc* genome (insertion at tRNA^gly^-gene copy 5) with bowtie2 using the default mode. Subsequent data analysis was performed using Seqmonk http://www.bioinformatics.babraham.ac.uk/projects/seqmonk/. A bin size of 1 kb was used. Graphs were plotted on GraphPad Prism for presentation.

### Expression and purification of ICEclc BisD protein

Expression constructs for full-length BisD and BisD(R95A) were prepared in a pLIBT7 derived plasmid by Golden-gate cloning with an N-terminal 6xHis-tag with an HRV-3C protease cleavage site (Table S1). Proteins were produced in *E. coli* BL21-Gold (DE3) grown in TB-medium at 37°C to an OD (600nm) of 1. Subsequently, the temperature of the culture was lowered to 24°C, and protein expression was initiated by adding IPTG to a concentration of 0.4 mM and was allowed to continue overnight, typically for 16 hours. Full-length His_6_-BisD protein was purified as follows. Cells were sonicated in lysis buffer A, which consisted of 500 mM NaCl, 50 mM Tris-HCl pH 7.5, 5 mM β-mercaptoethanol, 5 % (v/v) glycerol, 25 mM imidazole, and a protease inhibitor cocktail (Sigma). Following this, the supernatant was collected through centrifugation and loaded onto a 5-mL His-trap HP column (Cytivia). The column was then washed five times with the lysis buffer. The protein that remained bound to the column was then cleaved using HRV-3C Protease overnight, and the flowthrough sample was collected the next day. The sample was adjusted to a conductivity of 18 mS/cm by diluting with buffer B (50 mM Tris-HCl pH 7.5, 1 mM EDTA pH 7.5 and 2 mM β-mercaptoethanol) and loaded onto a Heparin column (GE Healthcare). The protein was eluted with a linear gradient of buffer B containing 1 M NaCl. The size of the eluted protein was verified on an SDS-PAGE. Peak fractions were then collected and directly loaded onto a Superdex 200 16/600 pg column (GE healthcare) that had been pre-equilibrated in 300 mM NaCl, 50 mM Tris-HCl pH 7.5, and 1 mM TCEP (tris(2-carboxyethyl)phosphine). Purity of the protein was verified by SDS-PAGE and the protein was aliquoted and flash frozen in liquid nitrogen, stored at –80°C.

### Isothermal titration calorimetry (ITC)

His_6_-BisD binding to ribonucleotides was measured by ITC in a MicroCal iTC200 (GE Healthcare Life Sciences) pre-cooled to 4°C. The buffer used for all measurements contained 150 mM NaCl, 50 mM Tris-HCl (pH 7.5), and 5 mM MgCl_2_. Both the measurement cell and injection syringe were rigorously cleaned with the buffer beforehand. The cell was loaded with 280 μL of a 60 μM protein solution, while the syringe was filled with 2 mM NTP solution (either ATP, CTP, GTP or UTP). The measurements were initiated after a 180-second delay, with the device set to a reference power of 5 μcal/sec, a stirring speed of 1000 rpm, and in ‘high feedback’ mode. The raw data, expressed in kcal/mol, were presented as a Wiseman plot, and regression curves were calculated using a 1:1 nucleotide-to-protein monomer binding model when applicable. Origin software (GE Healthcare) was employed for fitting the measurement results using the equation:

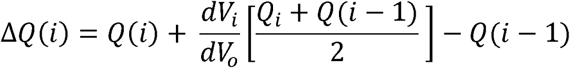

In this equation, *V_i_* is the ligand (nucleotide) volume, *V_0_* the cell volume, *Q*(*i*) the heat released from the *i^th^* injection which is in turn calculated using the following equation:

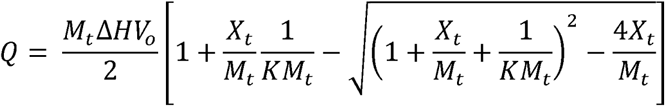

Here,*K* denotes the binding constant, Δ*H* the molar heat of ligand binding, *X_t_* the bulk nucleotide concentration, and *M_t_* is the bulk protein concentration (moles/liter) in *V_0_*.*K* and Δ*H* were estimated by Origin, and Δ*Q*(*i*) for each injection was calculated and compared to the measured heat. To refine the estimates of *K* and Δ*H*, standard Marquardt methods were applied, and iterative adjustments were made until no further improvement in the fit could be achieved.

### Measurement of NTP hydrolysis by Malachite Green colorimetric detection

The method for measuring NTP hydrolysis was adapted from the procedure outlined by (Antar et al., 2021). In brief, mixtures of NTP (2x conc.) with or without 40 nucleotide double-stranded DNA (“DNA_40_“) (2x conc.) and mixture of protein solutions (2x conc.) were prepared in reaction buffer (150 mM NaCl, 50 mM Tris/HCl pH 7.5, 5 mM MgCl_2_) on ice. Equal volumes of each solution were mixed, achieving a protein to ligand ratio of 1:1, and thoroughly mixed via pipetting. The resulting samples (containing 1 mM NTP, 1 μM DNA_40_, and 10 μM protein) were incubated at 25 °C for 1 hour, and phosphate standard curve were prepared in parallel. Post-incubation, the samples were diluted four-fold by the addition of 60 μL of water to stop the reaction. 20 μL of working reagent (Sigma) were then added before transferring to a flat-bottom 96-well plate. The plate was left to incubate for 30 minutes at 25 °C to allow for colour development, and then the absorbance was measure at at 620 nm wavelength. A standard curve of samples with increasing phosphate concentration was stained similarly and measured at 620 nm absorbance. Raw values were converted to rate values using the standard curve, and absolute rates were calculated by adjusting for the protein concentration. Finally, the mean values and standard deviations were calculated and graphically represented using GraphPad Prism software.

### Preparation of 40-nt double stranded DNA

40-nt double-stranded DNA was prepared by mixing solutions (at a concentration of 100 μM) of two complementary oligonucleotide strands (Table S2) in a 1:1 ratio. The resulting mixture was heated to 95 °C for 10 minutes and subsequently allowed to cool down slowly to 25 °C.

## Supplemental Files

Table S1 List of all used strains

Table S2 List of oligonucleotides

