## Supplementary tables and figures for "Orchestrated long-distance gene activation by a ParB-like BisD-CTP DNA clamp in low-frequency transfer competence development in *Pseudomonas putida*"

### Supplemental Files

Table S1 List of all used strains

Table S2 List of oligonucleotides

**Figure S1: Structural comparison** of AF3 prediction of ICEclc BisD dimer with the dimer of FlhC taken from a FlhDC co-crystal structure (PDB: 2avu). For simplicity, FlhD is omitted from the display. Shared structural features are indicated by dashed lines. Note that the C-terminal zinc finger domain of FlhD is not present in BisD.

**Figure S2: Fluorescent foci formation by mCherry-BisD under wild-type ICE*clc* induction conditions.** Micrographs show *P. putida* cells with integrated ICE*clc* grown on 3CBA (3-chlorobenzoate) until mid-(68 h) to late (72 h) stationary phase, showing the infrequent (3-5%) transfer competent cells with fluorescent foci for the variant with *mcherry-bisD* translational fusion replacement on the ICE, but homogenous fluorescence for the *bisC-mcherry* replacement. Foci formation correlates to cells expressing a single copy $P_{inR}$-*egfp* promoter (strain 7161/, but again only homogenous fluorescent for the BisC-mCherry expressing variant (strain 7163).

**Figure S3:** ChIP-Seq. Similar to Fig. 5 but showing the distribution profile of mCherry tagged BisD/C on the entire *Pseudomonas putida* ICE*clc* genome for a related experiment. Black curve represents the IP, and the orange represents the input from whole cell extract (WCE) DNA. All ChIP-Seq profiles were split into 1 kb bins.

**Figure S4:** ChIP-qPCR using anti-mCherry antibody. Enrichment was tested at three loci: at *bisS1*, *orf96323* (1 kb downstream *bisS1*) and (*clcA*) (80 kb downstream *bisS1*). Primer sequences used for qPCR are detailed in table S1.

**Figure S5**: Nucleotide binding by ITC. Same as in Fig.6 but for the BisD(R95A) mutant.

### Supplementary table 1: Strains and specifications

| strain numbe | Resistances | genotype | organism name | ICEclc or not | description | further description |
|---|---|---|---|---|---|---|
| 1291 | Rif | UWC1 | *Pseudomonas putida* UWC1 | no | wild-type strain | |
| 2737 | Rif | UWC1 ICEclc5 | *Pseudomonas putida* UWC1 | yes | one integrated ICEclc copy at glytRNA5 | |
| 2744 | Rif, Gm | UWCGC | *Pseudomonas putida* UWC1 | no | single copy Tn7 insertion of mcherry under Ptac control | |
| 5719 | Rif,Km,Tc | UWC1 Pint-echerry/PinR-egfp pME6032 | *Pseudomonas putida* UWC1 | no | single copy Tn5 insertion of Pint/PinR-double fluorescent reporter, clone 1 | plasmid pME6032 |
| 5724 | Rif,Km,Tc | UWC1 Pint-echerry/PinR-egfp pME97 | *Pseudomonas putida* UWC1 | no | single copy Tn5 insertion of Pint/PinR-double fluorescent reporter, clone 1 | plasmid pME6032 with cloned bisC under Ptac control |
| 5725 | Rif,Km,Tc | UWC1 Pint-echerry/PinR-egfp pME6032 | *Pseudomonas putida* UWC1 | no | single copy Tn5 insertion of Pint/PinR-double fluorescent reporter, clone 2 | plasmid pME6032 |
| 5730 | Rif,Km,Tc | UWC1 Pint-echerry/PinR-egfp pME97 | *Pseudomonas putida* UWC1 | no | single copy Tn5 insertion of Pint/PinR-double fluorescent reporter, clone 2 | plasmid pME6032 with cloned bisC under Ptac control |
| 5731 | Rif,Km,Tc | UWC1 Pint-echerry/PinR-egfp pME6032 | *Pseudomonas putida* UWC1 | no | single copy Tn5 insertion of Pint/PinR-double fluorescent reporter, clone 3 | plasmid pME6032 |
| 5736 | Rif,Km,Tc | UWC1 Pint-echerry/PinR-egfp pME97 | *Pseudomonas putida* UWC1 | no | single copy Tn5 insertion of Pint/PinR-double fluorescent reporter, clone 3 | plasmid pME6032 with cloned bisC under Ptac control |
| 5884 | Rif,Km,Tc | UWC1 Pint-echerry/PinR-egfp pMEparB | *Pseudomonas putida* UWC1 | no | single copy Tn5 insertion of Pint/PinR-double fluorescent reporter, clone 1 | plasmid pME6032 with cloned bisD under Ptac control |
| 5887 | Rif,Km,Tc | UWC1 Pint-echerry/PinR-egfp pMEparB | *Pseudomonas putida* UWC1 | no | single copy Tn5 insertion of Pint/PinR-double fluorescent reporter, clone 2 | plasmid pME6032 with cloned bisD under Ptac control |
| 5890 | Rif,Km,Tc | UWC1 Pint-echerry/PinR-egfp pMEparB | *Pseudomonas putida* UWC1 | no | single copy Tn5 insertion of Pint/PinR-double fluorescent reporter, clone 3 | plasmid pME6032 with cloned bisD under Ptac control |
| 6059 | Rif,Km,Tc | UWC1 Pint-echerry/PinR-egfp pMEparB97 | *Pseudomonas putida* UWC1 | no | single copy Tn5 insertion of Pint/PinR-double fluorescent reporter, clone 1 | plasmid pME6032 with cloned bisDC under Ptac control |
| 6060 | Rif,Km,Tc | UWC1 Pint-echerry/PinR-egfp pMEparB97 | *Pseudomonas putida* UWC1 | no | single copy Tn5 insertion of Pint/PinR-double fluorescent reporter, clone 2 | plasmid pME6032 with cloned bisDC under Ptac control |
| 6061 | Rif,Km,Tc | UWC1 Pint-echerry/PinR-egfp pMEparB97 | *Pseudomonas putida* UWC1 | no | single copy Tn5 insertion of Pint/PinR-double fluorescent reporter, clone 3 | plasmid pME6032 with cloned bisDC under Ptac control |
| 6317 | Tc | UWC1 ICEclc5 pMEbisR | *Pseudomonas putida* UWC1 | yes | plasmid pME6032 with bisR under Ptac control | |
| 7098 | Rif | UWC1 ICEclc5 mcherry-bisD | *Pseudomonas putida* UWC1 | yes, modified | mcherry with linker fused to bisD on ICEclc | |
| 7100 | Rif | UWC1 ICEclc5 bisC-mcherry | *Pseudomonas putida* UWC1 | yes, modified | bisC with linker fused to mcherry on ICEclc | |
| 7108 | Rif, Tc | UWC1 ICEclc5 mcherry-bisD pMEbisR | *Pseudomonas putida* UWC1 | yes, modified | mcherry with linker fused to bisD on ICEclc | plasmid pME6032 with cloned bisR under Ptac for population-wide induction of the ICE |
| 7110 | Rif, Tc | UWC1 ICEclc5 bisC-mcherry pMEbisR | *Pseudomonas putida* UWC1 | yes, modified | bisC with linker fused to mcherry on ICEclc | plasmid pME6032 with cloned bisR under Ptac for population-wide induction of the ICE |
| 7161 | Rif, Gm | UWC1 ICEclc5 mcherry-bisD miniTn7 PinR-gfp | *Pseudomonas putida* UWC1 | yes, modified | mcherry with linker fused to bisD on ICEclc | single copy Tn7 insertion of PinR-fluorescent reporter |
| 7163 | Rif, Gm | UWC1 ICEclc5 bisC-mcherry miniTn7 PinR-gfp | *Pseudomonas putida* UWC1 | yes, modified | bisC with linker fused to mcherry on ICEclc | single copy Tn7 insertion of PinR-fluorescent reporter |
| 7660 | Rif, Gm | UWC1 Tn7 Ptac-mcherry-bisD | *Pseudomonas putida* UWC1 | no | single copy Tn7 insertion of mcherry with linker fused to bisD under Ptac control | |
| 7680 | Rif, Tc | UWC1 pMEcheBDC | *Pseudomonas putida* UWC1 | no | plasmid pME6032 with mcherry with linker fused to bisD followed by bisC under Ptac control | |
| 7682 | Rif, Tc | UWC1 pMEcheB | *Pseudomonas putida* UWC1 | no | plasmid pME6032 with mcherry with linker fused to bisD until nt 1005 (only parB domain), under Ptac control | |
| 7684 | Rif, Tc | UWC1 ICEclc5 pMEcheBDC | *Pseudomonas putida* UWC1 | yes | plasmid pME6032 with mcherry with linker fused to bisD followed by bisC under Ptac control | |
| 7686 | Rif, Tc | UWC1 ICEclc5 pMEcheB | *Pseudomonas putida* UWC1 | yes | plasmid pME6032 with mcherry with linker fused to bisD until nt 1005 (only parB domain), under Ptac control | |
| 7692 | Rif,Km,Tc | UWC1 Pint-echerry/PinR-egfp pMEDC | *Pseudomonas putida* UWC1 | no | single copy Tn5 insertion of Pint/PinR-double fluorescent reporter, clone 1 | plasmid pME6032 with bisD starting from nt 1005 to end (only activator domain) and bisC under Ptac control |
| 7693 | Rif,Km,Tc | UWC1 Pint-echerry/PinR-egfp pMEDC | *Pseudomonas putida* UWC1 | no | single copy Tn5 insertion of Pint/PinR-double fluorescent reporter, clone 2 | |
| 7700 | Rif, Gm | UWC1 ICEclc5 Tn7::Ptac-mcherry-bisD | *Pseudomonas putida* UWC1 | yes | single copy Tn7 insertion of mcherry with linker fused to bisD under Ptac control | |
| 7717 | Rif, Gm, Km | UWC1 Tn7 Ptac-mcherry-bisD pBAMparS96323 | *Pseudomonas putida* UWC1 | no | single copy Tn7 insertion of mcherry with linker fused to bisD under Ptac control | single copy Tn5 insertion of bisS-3 site, clone 1 |
| 7718 | Rif, Gm, Km | UWC1 Tn7 Ptac-mcherry-bisD pBAMparS96323 | *Pseudomonas putida* UWC1 | no | single copy Tn7 insertion of mcherry with linker fused to bisD under Ptac control | single copy Tn5 insertion of bisS-3 site, clone 2 |
| 7719 | Rif, Gm, Km | UWC1 Tn7 Ptac-mcherry-bisD pBAMparS96323 | *Pseudomonas putida* UWC1 | no | single copy Tn7 insertion of mcherry with linker fused to bisD under Ptac control | single copy Tn5 insertion of bisS-3 site, clone 3 |
| 7720 | Rif, Gm, Km | UWC1 Tn7 Ptac-mcherry-bisD pBAMparS9596 | *Pseudomonas putida* UWC1 | no | single copy Tn7 insertion of mcherry with linker fused to bisD under Ptac control | single copy Tn5 insertion of bisS-1 site, clone 1 |
| 7721 | Rif, Gm, Km | UWC1 Tn7 Ptac-mcherry-bisD pBAMparS9596 | *Pseudomonas putida* UWC1 | no | single copy Tn7 insertion of mcherry with linker fused to bisD under Ptac control | single copy Tn5 insertion of bisS-1 site, clone 2 |
| 7722 | Rif, Gm, Km | UWC1 Tn7 Ptac-mcherry-bisD pBAMparS9596 | *Pseudomonas putida* UWC1 | no | single copy Tn7 insertion of mcherry with linker fused to bisD under Ptac control | single copy Tn5 insertion of bisS-1 site, clone 3 |
| 7723 | Rif, Gm, Km | UWC1 Tn7 Ptac-mcherry-bisD pBAMparSssb | *Pseudomonas putida* UWC1 | no | single copy Tn7 insertion of mcherry with linker fused to bisD under Ptac control | single copy Tn5 insertion of bisS-2 site, clone 1 |
| 7724 | Rif, Gm, Km | UWC1 Tn7 Ptac-mcherry-bisD pBAMparSssb | *Pseudomonas putida* UWC1 | no | single copy Tn7 insertion of mcherry with linker fused to bisD under Ptac control | single copy Tn5 insertion of bisS-2 site, clone 2 |
| 7725 | Rif, Gm, Km | UWC1 Tn7 Ptac-mcherry-bisD pBAMparSssb | *Pseudomonas putida* UWC1 | no | single copy Tn7 insertion of mcherry with linker fused to bisD under Ptac control | single copy Tn5 insertion of bisS-2 site, clone 3 |
| 7726 | Rif, Tc, Km | UWC1 pMEcheBDC pBAMparSssb | *Pseudomonas putida* UWC1 | no | single copy Tn5 insertion of bisS-2 site, clone 1 | plasmid pME6032 with mcherry with linker fused to bisD followed by bisC under Ptac control |
| 7727 | Rif, Tc, Km | UWC1 pMEcheBDC pBAMparSssb | *Pseudomonas putida* UWC1 | no | single copy Tn5 insertion of bisS-2 site, clone 2 | plasmid pME6032 with mcherry with linker fused to bisD followed by bisC under Ptac control |
| 7728 | Rif, Gm, Km | UWC1 Tn7::Ptac-mcherry-bisD pBAMgfp | *Pseudomonas putida* UWC1 | no | single copy Tn7 insertion of mcherry with linker fused to bisD under Ptac control | single copy mini-Tn5 insertion of egfp without promoter |
| 7850 | Rif, Tc | ICEclc5 cherry-bisDnolinker D21 pMEbisR | *Pseudomonas putida* UWC1 | yes, modified | native bisD replaced by mcherry fused to bisD that has no linker between the parB and | plasmid pME6032 with cloned bisR funder Ptac for population-wide induction of the ICE |
| 7865 | Rif, Gm | ICEclc5 cherry-bisDnolinker D21 miniTn7::PinR-gfp | *Pseudomonas putida* UWC1 | yes, modified | native bisD replaced by mcherry fused to bisD that has no linker between the parB and | single copy Tn7 insertion of PinR-fluorescent reporter |
| pSG4830 | Km | pLIBT7 Kan His6-3C-BisD | *Escherichia coli* BL21-Gold (DE | no | cloning of bisD in pLIBT7 for overexpression with N-terminal His6 tag | |
| pSG7787 | Km | pLIBT7 Kan His6-3C-BisD(R95A) | *Escherichia coli* BL21-Gold (DE | no | cloning of bisD(R95A) in pLIBT7 for overexpression with N-terminal His6 tag | |

Supplementary Table 2: Primers used in the study

| Identifier | Sequence 5'-3' | Locus |
|---|---|---|
| STP905 | CATGGCGTTTGATGGAGCTA | Forward *bisS-1* for qPCR |
| STP906 | GGAAGGCAGTCAGTCAACAA | Reverse *bisS-1* for qPCR |
| STP907 | GATGATGCTCAACGAGGATG | Forward *orf96323* for qPCR |
| STP908 | GAGACTCAACCAGCGTGTCA | Reverse *orf96323* for qPCR |
| STP909 | CACGCTCGATTACGAAGTCC | Forward *clcA* for qPCR |
| STP910 | ACTACTTCGAAGGGGGCAAA | Reverse *clcA* for qPCR |
| STP747 | GTTTGATGGAGCTATCCCCTGGGGATAGCTCGTGCCTCGG | Forward *bisS-1* [40] |
| STP748 | CCGAGGCACGAGCTATCCCCAGGGGATAGCTCCATCAAAC | Reverse *bisS-1* [40] |
| STP532 | ATACTTCCGGGGGCCCAGCTCGACCCGGAAGTATATTTAA | Forward *random* [40] |
| STP533 | TTAAATATACTTCCGGGTCGAGCTGGGCCCCCGGAAGTAT | Reverse *random* [40] |
| 220207 | acaggaaacagaattc GAGCTC AAGGAGGA TGCGGCCATGGCTGATA | Forward primer for the amplification of echerry-bisD fragment. The primer contains a SacI restriction site and RBS (AAGGAGGA). The primer overlaps with the Tn7 Ptac plasmid. |
| 220208 | aggtaccgggcccaagctt CTCGAG TCAGGTCCCAGAACCGAC | Reverse primer for the amplification of echerry-bisD fragment. The primer contains a XhoI restriction site and overlaps with the Tn7 Ptac plasmid. |
| 220212 | GTCGGTTCTGGGACCTGA CTCGAG AAGGAGGA TCACACCATGAGCACGC | Forward primer for the amplification of orf94175 (SSB) from ICEclc. The primer contains a XhoI restriction site, a RBS (AAGGAGGA) and overlaps with the bisD gene cloned in the Tn7 mcherry-bisD vector. |
| 200305 | ATGGTTTCCAAGGGCGAGGA | Amplification of mcherry from pBam-Cherry-link (for) |
| 200306 | AAGCTTGGTGGCCGACCGGT | Amplification of mcherry from pBam-Cherry-link (rev) |
| 200307 | AAGCTTCCGGAAAATTCGAA | Amplification of mcherry from pBam-link-Cherry (for) |
| 200308 | TTATTTGTACAGCTCATCCAT | Amplification of mcherry from pBam-link-Cherry (rev) |
| 200325 | AGGGATAACAGGGTAATCTGAATTCCCCACGTTGTCGTCCTATTA | Amplification upstream flanking region of bisD (for) |
| 200326 | CTCCTCGCCCTTGGAAAC**CAT**A**TCA**GCCATGGCCGCATTCCT | Amplification upstreamflanking region of bisD (rev) |
| 200327 | CACCGGTCGGCCACCAAGCTT**ATG**GCTGATATGAGGTCCCA | Amplification downstream flanking region of bisD (for) |
| 200328 | GAAGCTTGCATGCCTGCAGGTCGACGATAGTGGATGGTGCTGATT | Amplification downstream flanking region of bisD (rev) |
| 200329 | AGGGATAACAGGGTAATCTGAATTCGCCGGTGGCCTCTATCCCAT | Amplification upstream flanking region of bisC (for) |
| 200330 | GTTCGAATTTTCCGGAAGCTTACCCAATCCTTGATCGAT | Amplification upstreamflanking region of bisC (rev) |
| 200331 | ATGGATGAGCTGTACAAA**TAA**GTTATGGCCGTGGATGACT | Amplification downstream flanking region of bisC (for) |
| 200332 | GAAGCTTGCATGCCTGCAGGTCGACCCTGCTGCAATGCCACCAT | Amplification downstream flanking region of bisC (rev) |
| pME6032 EcorI (5') | AACAATTTCACACAGGAAACA**GAATTC** | Cloning of bisD in pME6032 |
| pME6032 XhoI (3') | CTGATCCGCTAGTCCGAGGC**CTCGAG** | Cloning of bisD in pME6032 |
| Cherrylink.F | **ATG**GTTTCCAAGGGCGAGGA | Cloning of mCherry-link-bisD in pME6032 |
| linkCherry.R | **TCATTA**TTTGTACAGCTCATCCAT | Cloning of mCherry-link-bisD in pME6032 |
| startofparB.f | **ATG**GCTGATATGAGGTCCCA | Cloning of mCherry-link-bisD in pME6032 |
| endofbisD.r | **TCA**GGTCCCAGAACCGACA | Cloning of mCherry-link-bisD in pME6032 |
| parB1005.r | **TTATCA**TGGACGTGGTGCAGGCGTGAT | Amplification of bisD until nt 1005 - only parB domain |
| bisD1005.f | **ATG**GGCCATGCAGCGGCCGCGCCTA | Amplification of bisD until nt 1005 - only parB domain |
| endofbisC.r | **TTATCA**ACCCAATCCTTGATCGAT | Primer end of bisC |
| cheBDC.f | AACAATTTCACACAGGAAACA**GAATTC**GGAA**GGAG***ACGGAGC* **ATG**GTTTCCAAGGGCGAGGA | Cloning of mCherry-link-bisD-bisC in pME6032 |
| cheBDC.r | CTGATCCGCTAGTCCGAGGC**CTCGAGTTATCA**ACCCAATCCTTGATCGAT | Cloning of mCherry-link-bisD-bisC in pME6032 |
| BDCche.f | AACAATTTCACACAGGAAACA**GAATTC**GGAA**GGAG***ACGGAGC* **ATG**GCTGATATGAGGTCCCA | Cloning of mCherry-link-bisD-bisC in pME6032 |
| BDCche.r | CTGATCCGCTAGTCCGAGGC**CTCGAGTCATTA**TTTGTACAGCTCATCCAT | Cloning of mCherry-link-bisD-bisC in pME6032 |
| parB1005.r | CTGATCCGCTAGTCCGAGGC**CTCGAGTTATCA**TGGACGTGGTGCAGGCGTGAT | Cloning of mCherry-link-bisD (parA domain) |
| bisD1005.f | AACAATTTCACACAGGAAACA**GAATTC**GGAA**GGAG***ACGGAGC* **ATG**GGCCATGCAGCGGCCGCGCCT | Cloning of mCherry-link-bisD (parA domain) |
| parS96323.F | TTCGAGGCATGCCTGCAGCCCGTTATCCACAGGAGGACAT | Amplification of bisS-3 site |
| parS96323.R | AATCAGAATTCGAGCTCGCCCCGACTACTACGTACAGTA | Amplification of bisS-3 site |
| parSssb.F | TTCGAGGCATGCCTGCAGCCCTGGCGAGTGGCCGGAGGTA | Amplification of bisS-2 site |
| parSssb.R | AATCAGAATTCGAGCTCGCCCGACGAAGCGCAGGAAGTGA | Amplification of bisS-2 site |
| parS95-96.F | TTCGAGGCATGCCTGCAGCCCGTCAACAAACCGAAATGAAT | Amplification of bisS-1 site |
| parS95-96.R | AATCAGAATTCGAGCTCGCCCGCTGATCGTTACGAGCGGCAT | Amplification of bisS-1 site |
| parSssb/96323.R | AATCAGAATTCGAGCTCGCCCTGGCGAGTGGCCGGAGGTA | Amplification of bisS-1 and bisS-2 sites on single fragment |
| parS95-96.R2 | AATCAGAATTCGAGCTCGCCCGAGGCACCACGCAATGTGGTT | Amplification of bisS-1 and bisS-2 sites on single fragment |
| DelParBUP.F | AGGGATAACAGGGTAATCTGAATTC CATGCAGGTTGTATCCATCAT | Amplification of upstream fragment for parB-domain deletion |
| DelParBUP.R | CATATCAGC**CAT**GGCCGCAT | Amplification of upstream fragment for parB-domain deletion |
| DelParBDW.F | GAATGCGGCC**ATG**GCTGATATG GGCCATGCAGCGGCCGCGCCTA | Amplification of downstream fragment for parB-domain deletion |
| DelParBDW.R | GAAGCTTGCATGCCTGCAGGTCGAC CTTCCAGCGCCGCCACAGTT | Amplification of downstream fragment for parB-domain deletion |
| DelParBlinkDW.F | GAATGCGGCC**ATG**GCTGATATG GCCAGCCCTACGGCAGGGGT | Amplification of bisD linker region |
| SplitBisDUP.F | AGGGATAACAGGGTAATCTGAATTC TCCCAGGACATGGCTGGCAA | Amplification of upstream fragment for deletion of bisD linker region |
| SplitBisDUP.R | **CAT**ATGTTTTT**CCTCCTT**AATTATCACTA TGGACGTGGTGCAGGCGTGAT | Amplification of upstream fragment for deletion of bisD linker region |
| SplitBisDDW.F | TAGTGATAATT**AAGGAGG**AAAAACAT**ATG** GGCCATGCAGCGGCCGCGCCTA | Amplification of downstream fragment for deletion of bisD linker region |
| SplitBisDDW.R | GAAGCTTGCATGCCTGCAGGTCGAC CTTCCAGCGCCGCCACAGTT | Amplification of downstream fragment for deletion of bisD linker region |
| SplitBisDlink.F | TAGTGATAATT**AAGGAGG**AAAAACAT**ATG** GCCAGCCCTACGGCAGGGGT | Amplification of bisD linker region |
| DellinkUP.R | CGGGTCGCTGACCAACGCA | Amplification of bisD linker region |
| DellinkDW.F | CACCGTGCGTTGGTCAGCGACCCG GCCAGCCCTACGGCAGGGGT | Amplification of bisD linker region |

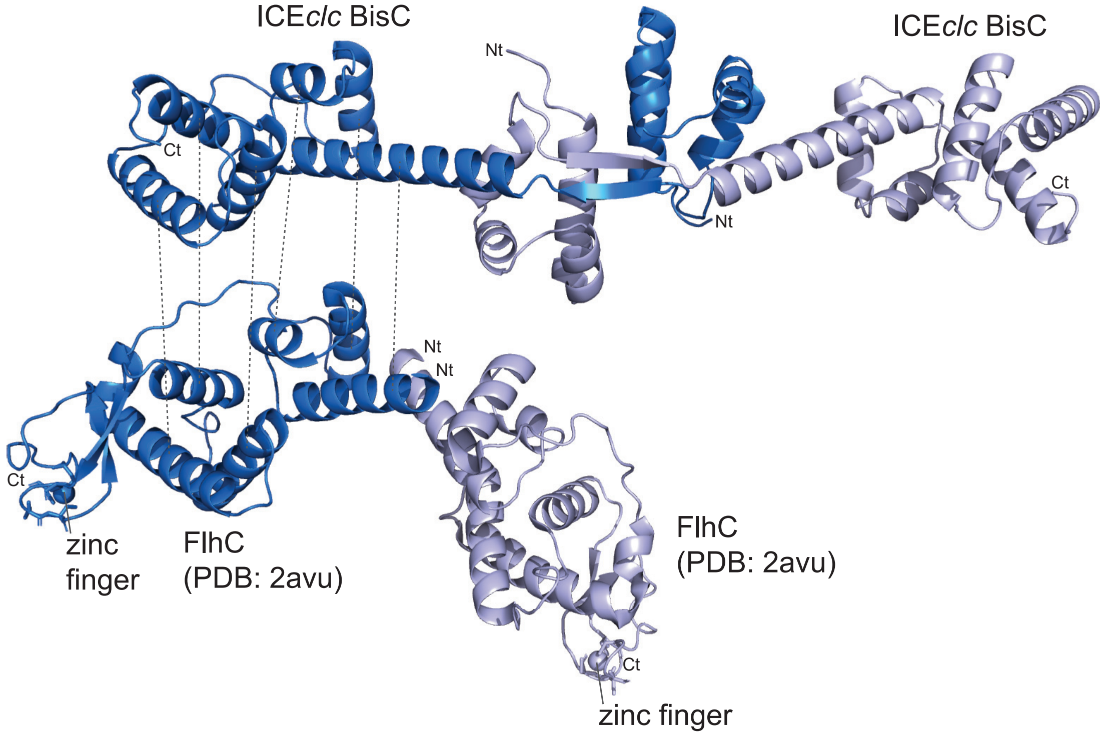
ICEclc BisC
ICEclc BisC
Nt
Ct
Nt
Ct
Nt
Nt
Ct
zinc finger
FlhC (PDB: 2avu)
FlhC (PDB: 2avu)
Ct
zinc finger

Fig. S1

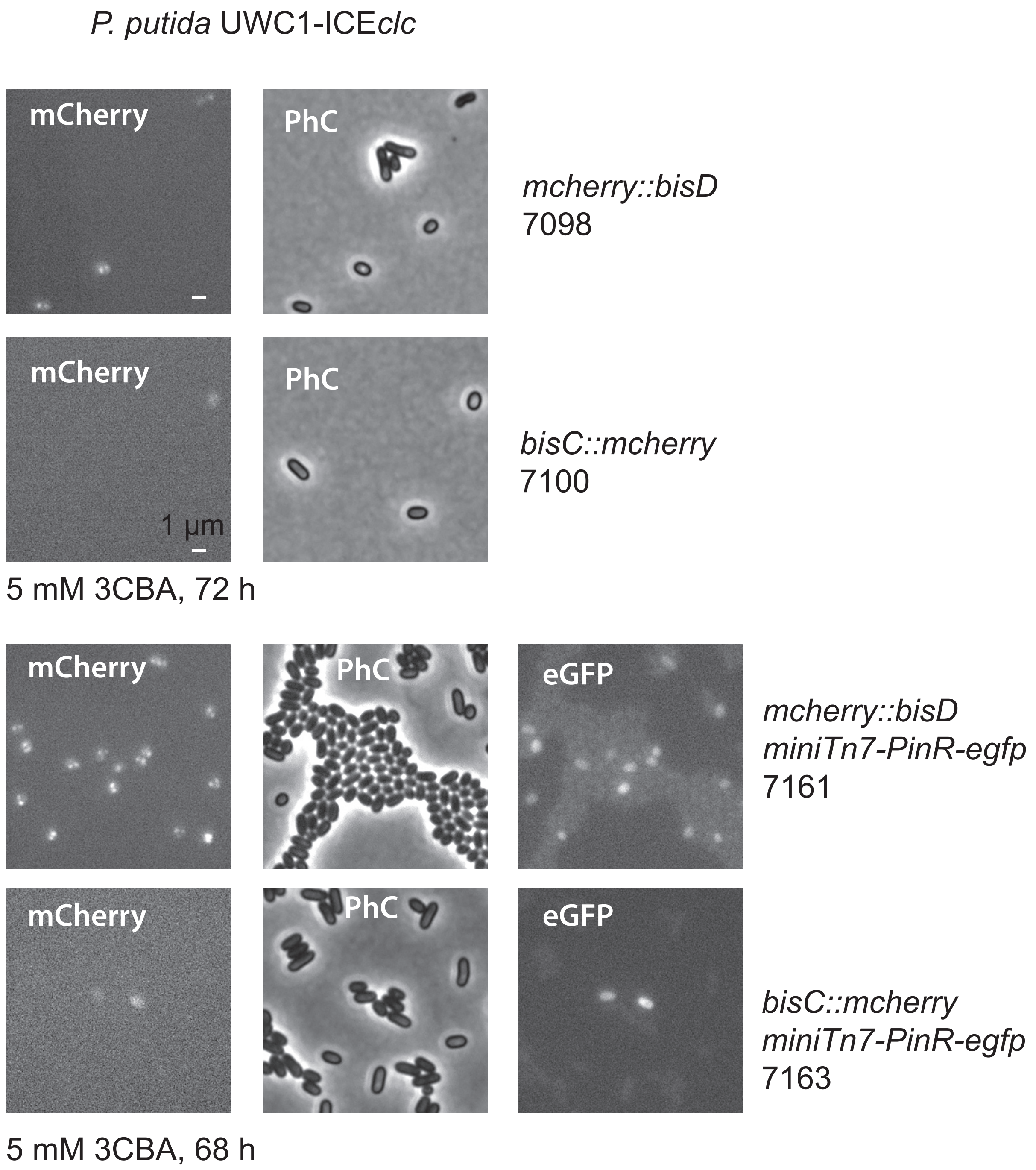
P. putida UWC1-ICEclc
mCherry
PhC
mcherry::bisD
7098
mCherry
PhC
1 µm
bisC::mcherry
7100
5 mM 3CBA, 72 h
mCherry
PhC
eGFP
mcherry::bisD
miniTn7-PinR-egfp
7161
mCherry
PhC
eGFP
bisC::mcherry
miniTn7-PinR-egfp
7163
5 mM 3CBA, 68 h

Fig. S2

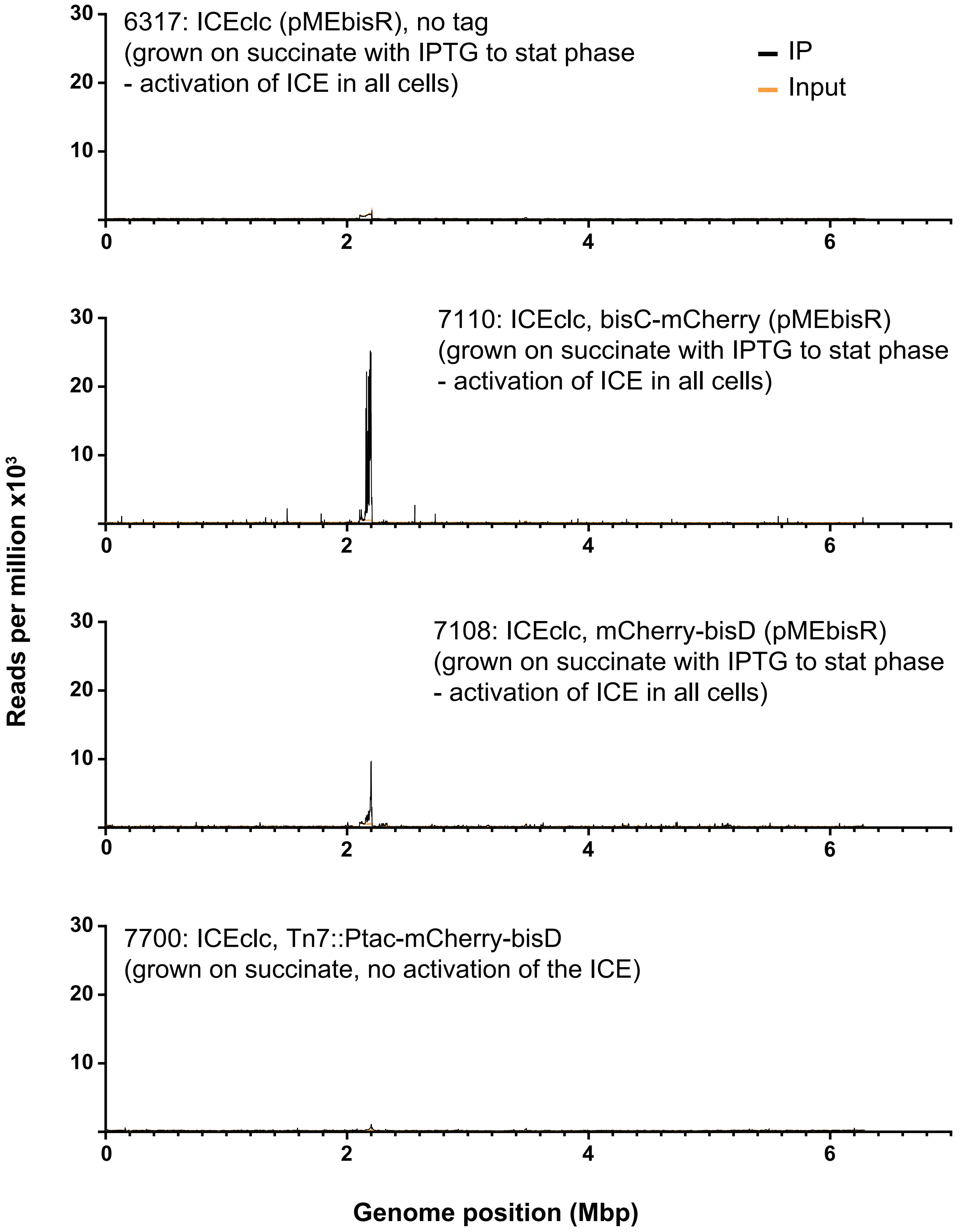
6317: ICEclc (pMEbisR), no tag
(grown on succinate with IPTG to stat phase
- activation of ICE in all cells)
IP
Input
7110: ICEclc, bisC-mCherry (pMEbisR)
(grown on succinate with IPTG to stat phase
- activation of ICE in all cells)
7108: ICEclc, mCherry-bisD (pMEbisR)
(grown on succinate with IPTG to stat phase
- activation of ICE in all cells)
7700: ICEclc, Tn7::Ptac-mCherry-bisD
(grown on succinate, no activation of the ICE)
Reads per million x10³
Genome position (Mbp)
0
2
4
6
10
20
30

Fig. S3

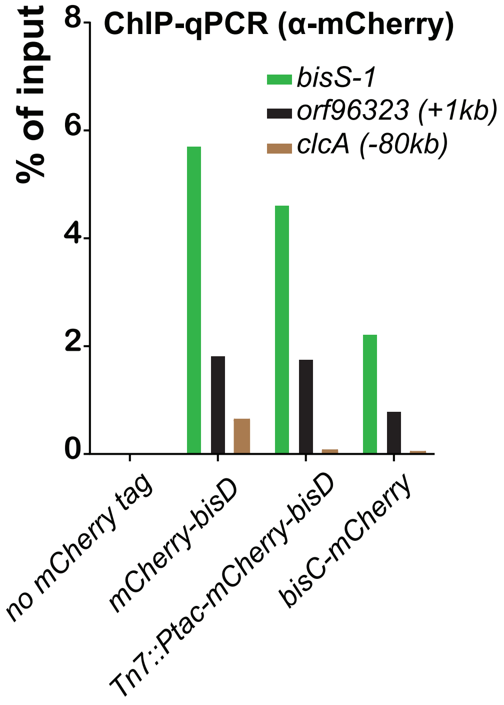
ChIP-qPCR (α-mCherry)
% of input
8
6
4
2
0
bisS-1
orf96323 (+1kb)
clcA (-80kb)
no mCherry tag
mCherry-bisD
Tn7::Ptac-mCherry-bisD
bisC-mCherry

Fig. S4

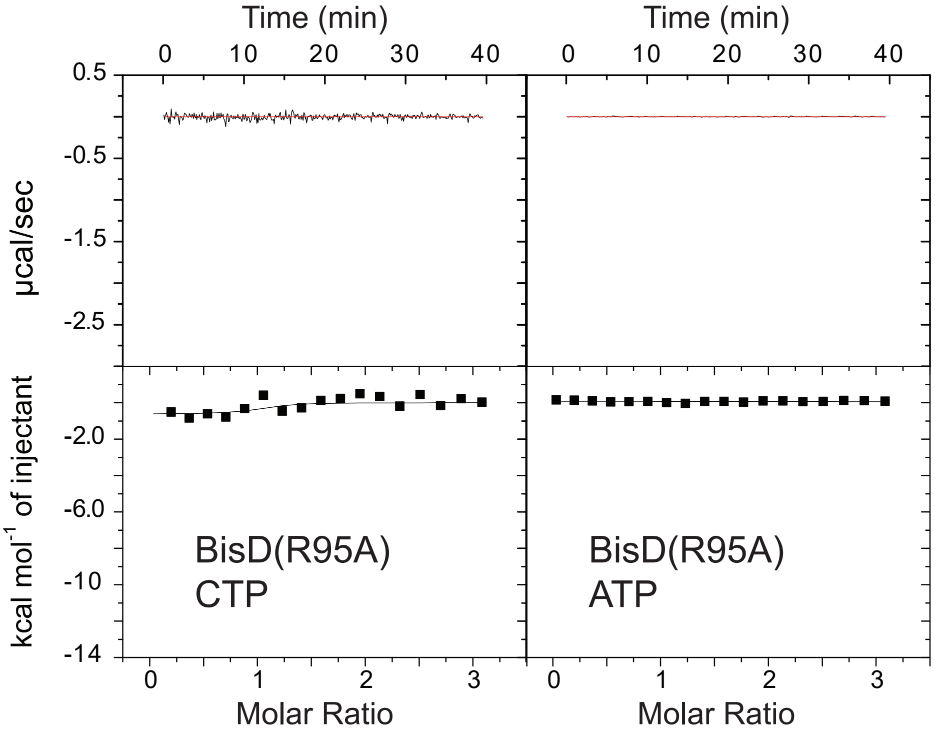
Time (min)
0 10 20 30 40
0.5
-0.5
-1.5
-2.5
µcal/sec
kcal mol-1 of injectant
-2.0
-6.0
-10
-14
0 1 2 3
Molar Ratio
BisD(R95A)
CTP
BisD(R95A)
ATP

Fig. S5
